# Cytoplasmic mRNA decay represses RNA polymerase II transcription during early apoptosis

**DOI:** 10.1101/2020.04.09.034694

**Authors:** Christopher Duncan-Lewis, Ella Hartenian, Valeria King, Britt A. Glaunsinger

## Abstract

RNA abundance is generally sensitive to perturbations in decay and synthesis rates, but crosstalk between RNA polymerase II transcription and cytoplasmic mRNA degradation often leads to compensatory changes in gene expression. Here, we reveal that widespread mRNA decay during early apoptosis represses RNAPII transcription, indicative of positive (rather than compensatory) feedback. This repression requires active cytoplasmic mRNA degradation, which leads to impaired recruitment of components of the transcription preinitiation complex to promoter DNA. Importin α/β- mediated nuclear import is critical for this feedback signaling, suggesting that proteins translocating between the cytoplasm and nucleus connect mRNA decay to transcription. We also show that an analogous pathway activated by viral nucleases similarly depends on nuclear protein import. Collectively, these data demonstrate that accelerated mRNA decay leads to the repression of mRNA transcription, thereby amplifying the shutdown of gene expression. This highlights a conserved gene regulatory mechanism by which cells respond to threats.

**IMPACT STATEMENT:** Human cells respond to cytoplasmic mRNA depletion during early apoptosis by inhibiting RNA polymerase II transcription, thereby magnifying the gene expression shutdown during stress.

## INTRODUCTION

Gene expression is often depicted as a unidirectional flow of discrete stages: DNA is first transcribed by RNA polymerase II (RNAPII) into messenger RNA (mRNA), which is processed and exported to the cytoplasm where it is translated and then degraded. However, there is a growing body of work that reveals complex cross talk between the seemingly distal steps of transcription and decay. For example, the yeast Ccr4-Not deadenylase complex, which instigates basal mRNA decay by removing the poly(A) tails of mRNAs (Tucker et al., 2001), was originally characterized as a transcriptional regulator (Collart & Stmhp, 1994; Denis, 1984). Other components of transcription such as RNAPII subunits and gene promoter elements have been linked to cytoplasmic mRNA decay (Bregman et al., 2011; Lotan et al., 2005; Lotan, Goler-Baron, Duek, Haimovich, & Choder, 2007), while the activity of cytoplasmic mRNA degradation machinery such as the cytoplasmic 5’-3’ RNA exonuclease Xrn1 can influence the transcriptional response (Haimovich et al., 2013; Sun et al., 2012).

The above findings collectively support a model in which cells engage a buffering response to reduce transcription when mRNA decay is slowed, or reduce mRNA decay when transcription is slowed to preserve the steady state mRNA pool (Haimovich et al., 2013; Hartenian & Glaunsinger, 2019). While much of this research has been performed in yeast, the buffering model is also supported by studies in mouse and human cells (Helenius et al., 2011; Singh et al., 2019). In addition to bulk changes to the mRNA pool, compensatory responses can also occur at the individual gene level to buffer against aberrant transcript degradation. Termed “nonsense-induced transcription compensation” (NITC; Wilkinson, 2019), this occurs when nonsense-mediated mRNA decay leads to transcriptional upregulation of genes with some sequence homology to the aberrant transcript (El-Brolosy et al., 2019; Ma et al., 2019).

A theme that unites much of the research linking mRNA decay to transcription is homeostasis; perturbations in mRNA stability are met with an opposite transcriptional response in order to maintain stable mRNA transcript levels. However, there are cellular contexts in which homeostasis is not beneficial, for example during viral infection. Many viruses induce widespread host mRNA decay (Narayanan & Makino, 2013) and co-opt the host transcriptional machinery (Harwig, Landick, & Berkhout, 2017) in order to express viral genes. Indeed, infection with mRNA decay-inducing herpesviruses or expression of broad-acting viral ribonucleases in mammalian cells causes RNAPII transcriptional repression in a manner linked to accelerated mRNA decay (Abernathy, Gilbertson, Alla, & Glaunsinger, 2015; Hartenian, Gilbertson, Federspiel, Cristea, & Glaunsinger, 2020). It is possible that this type of positive feedback represents a protective cellular shutdown response, perhaps akin to the translational shutdown mechanisms that occur upon pathogen sensing (D. Walsh, Mathews, & Mohr, 2013). A central question, however, is whether transcriptional inhibition upon mRNA decay is restricted to infection contexts, or whether it is also engaged upon other types of stimuli.

The best-defined stimulus known to broadly accelerate cytoplasmic mRNA decay outside of viral infection is induction of apoptosis. Overall levels of poly(A) RNA are reduced rapidly after the induction of extrinsic apoptosis via accelerated degradation from the 3’ ends of transcripts (Thomas et al., 2015). The onset of accelerated mRNA decay occurs coincidentally with mitochondrial outer membrane depolarization (MOMP) and requires release of the mitochondrial exonuclease polyribonucleotide nucleotidyltransferase 1 (PNPT1) into the cytoplasm. PNPT1 then coordinates with other 3’ end decay machinery such as DIS3L2 and terminal uridylyltransferases (TUTases; Liu et al., 2018; Thomas et al., 2015). Notably, mRNA decay occurs before other hallmarks of apoptosis including phosphatidylserine (PS) externalization and DNA fragmentation, but likely potentiates apoptosis by reducing the expression of unstable anti-apoptotic proteins such as MCL1 (Thomas et al., 2015).

Here, we used early apoptosis as a model to study the impact of accelerated cytoplasmic mRNA decay on transcription. We reveal that under conditions of increased mRNA decay, there is a coincident decrease in RNAPII transcription, indicative of positive feedback between mRNA synthesis and degradation. Using decay factor depletion experiments, we demonstrate that mRNA decay is required for the transcriptional decrease and further show that transcriptional repression is associated with reduced RNAPII polymerase occupancy on promoters. This phenotype requires ongoing nuclear-cytoplasmic protein transport, suggesting that protein trafficking may provide the signal linking cytoplasmic decay to transcription. Collectively, our findings elucidate a distinct gene regulatory mechanism by which cells sense and respond to threats.

## RESULTS

### mRNA decay during early apoptosis is accompanied by reduced synthesis of RNAPII transcripts

To induce widespread cytoplasmic mRNA decay, we initiated rapid extrinsic apoptosis in HCT116 colon carcinoma cells by treating them with TNF-related apoptosis inducing ligand (TRAIL). TRAIL treatment causes a well-characterized progression of apoptotic events including caspase cleavage and mitochondrial outer membrane permeabilization or “MOMP” (Albeck et al., 2008; Thomas et al., 2015). It is MOMP that stimulates mRNA decay in response to an apoptosis-inducing ligand (Figure 1A), partly by releasing the mitochondrial 3’-5’ RNA exonuclease PNPT1 into the cytoplasm (Liu et al., 2018; Thomas et al., 2015). A time course experiment in which cells were treated with 100 ng/mL TRAIL for increasing 30 min increments showed activation of caspase 8 (CASP8) and caspase 3 (CASP3) by 1.5 hr (Figure 1B), as measured by disappearance of the full-length zymogen upon cleavage (Kim et al., 2000; Thomas et al., 2015). In agreement with Liu et al. (2018), RT-qPCR performed on total RNA revealed a coincident decrease in the mRNA levels of several housekeeping genes (*ACTB*, *GAPDH*, *EEF1A*, *PPIA, CHMP2A, DDX6, RPB2, and RPLP0*) beginning 1.5 hr after TRAIL was applied (Figure 1C, Supplementary Figure 1A). Fold changes were calculated in reference to 18S rRNA, which has been shown to be stable during early apoptosis (Houge et al., 1995; Thomas et al., 2015). As expected, this decrease was specific to RNAPII transcripts, as the RNA polymerase III (RNAPIII)-transcribed ncRNAs *U6*, *7SK, 7SL, and 5S* did not show a similarly progressive decrease. The *U6* transcript was instead upregulated, possibly suggesting post-transcriptional regulation of *U6* as alluded to in a previous study (Noonberg, Scott, & Benz, 1996). These data confirm that mRNA depletion occurs by 1.5-2 hr during TRAIL-induced apoptosis.

**Figure 1.**
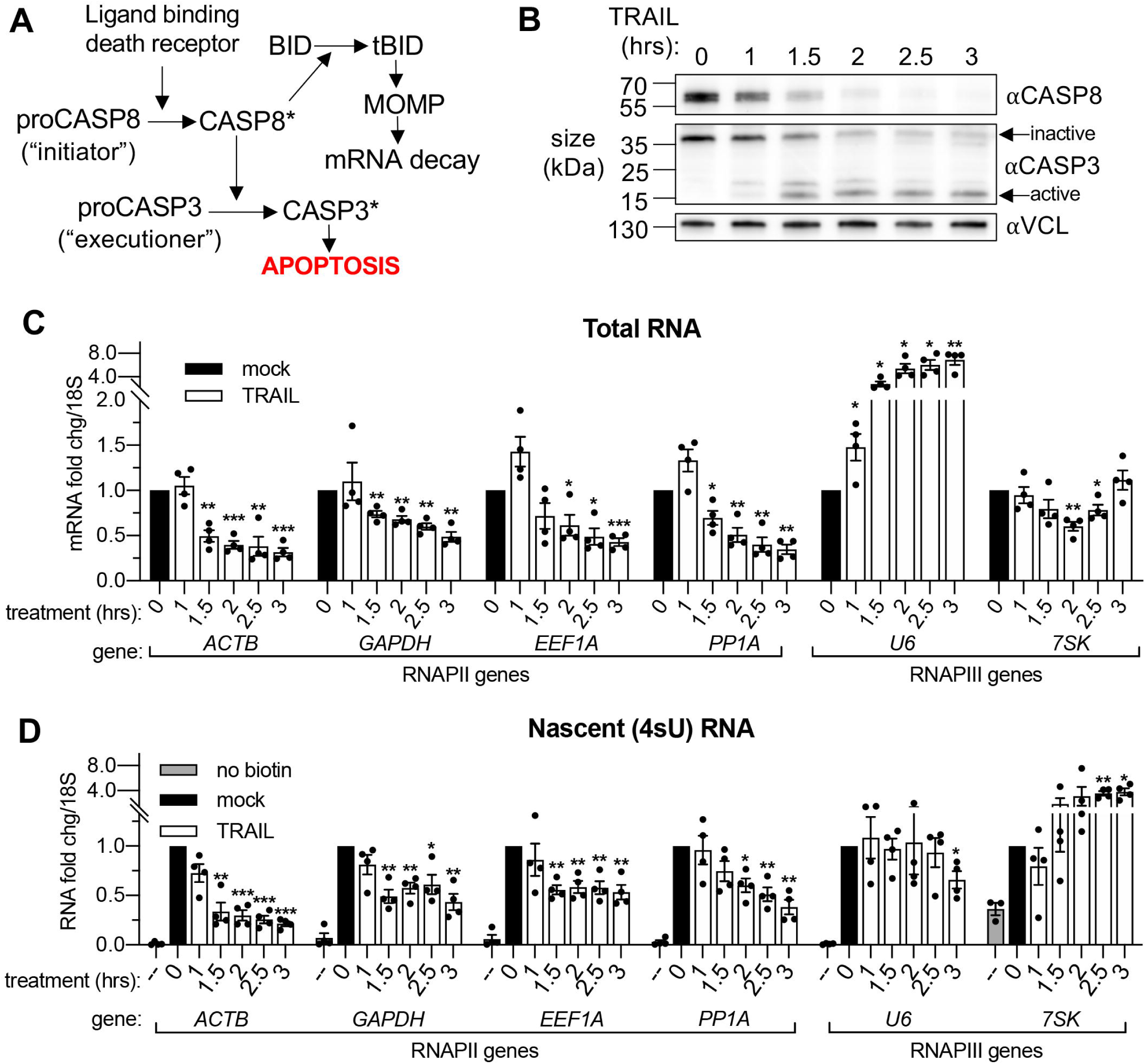
mRNA decay during early apoptosis is accompanied by reduced synthesis of RNAPII transcripts. (A) Schematic representation of how the extrinsic apoptotic pathway accelerates mRNA decay. (B) Western blot of HCT116 lysates showing the depletion of full-length caspase 8 (CASP8) and caspase 3 (CASP3) over a time course of 100 ng/μL TRAIL treatment. Vinculin (VCL) serves as a loading control. Blot representative of those from 4 biological replicates. (C, D) RT-qPCR quantification of total (C) and nascent 4sU pulse-labeled (D) RNA at the indicated times post TRAIL treatment of HCT116 cells (n = 4). Also see Supplementary Figure 1. No biotin control quantifies RNA not conjugated to biotin that is pulled down with strepdavidin selection beads. Fold changes were calculated from Ct values normalized to 18S rRNA in reference to mock treated cells. Graphs display mean +/- SEM with individual biological replicates represented as dots. Statistically significant deviation from a null hypothesis of 1 was determined using one sample t test; *p<0.05, **p<0.01, ***p<0.001.

To monitor whether apoptosis also influenced transcription, we pulse labeled the cells with 4-thiouridine (4sU) for 20 min at the end of each TRAIL treatment. 4sU is incorporated into actively transcribing RNA and can be subsequently coupled to HPDP-biotin and purified over streptavidin beads, then quantified by RT-qPCR to measure nascent transcript levels (Dölken, 2013). 4sU-labeled RNA levels were also normalized to 18S rRNA, which was produced at a constant level in the presence and absence of TRAIL (Supplementary Figure 1B). In addition to a reduction in steady state mRNA abundance, TRAIL treatment caused a decrease in RNAPII-driven mRNA production, while RNAPIII transcription was largely either unaffected or enhanced (Figure 1D, Supplementary Figure 1C). Thus, TRAIL triggers mRNA decay and decreases nascent mRNA production in HCT116 cells but does not negatively impact RNAPIII transcript abundance or production.

### RNAPII transcription is globally repressed during early apoptosis

z-VAD-fmk (zVAD), a pan-caspase inhibitor, was used to confirm that TRAIL-induced mRNA decay and transcriptional arrest were associated with apoptosis and not due to an off-target effect of TRAIL. HCT116 cells were pre-treated with 40 μM zVAD or equal volume of vehicle (DMSO) for 1 hr prior to TRAIL treatment. The effectiveness of zVAD treatment was confirmed by showing it blocked the cleavage of the CASP8 target BID and blocked the degradation of the CASP3 substrate PARP1 (Supplementary Figure 2A). The decreases in total and nascent 4sU-labeled mRNA abundance upon TRAIL treatment were rescued in the presence of zVAD (Figure 2A-B), confirming the role of canonical apoptotic signaling in both phenotypes.

**Figure 2.**
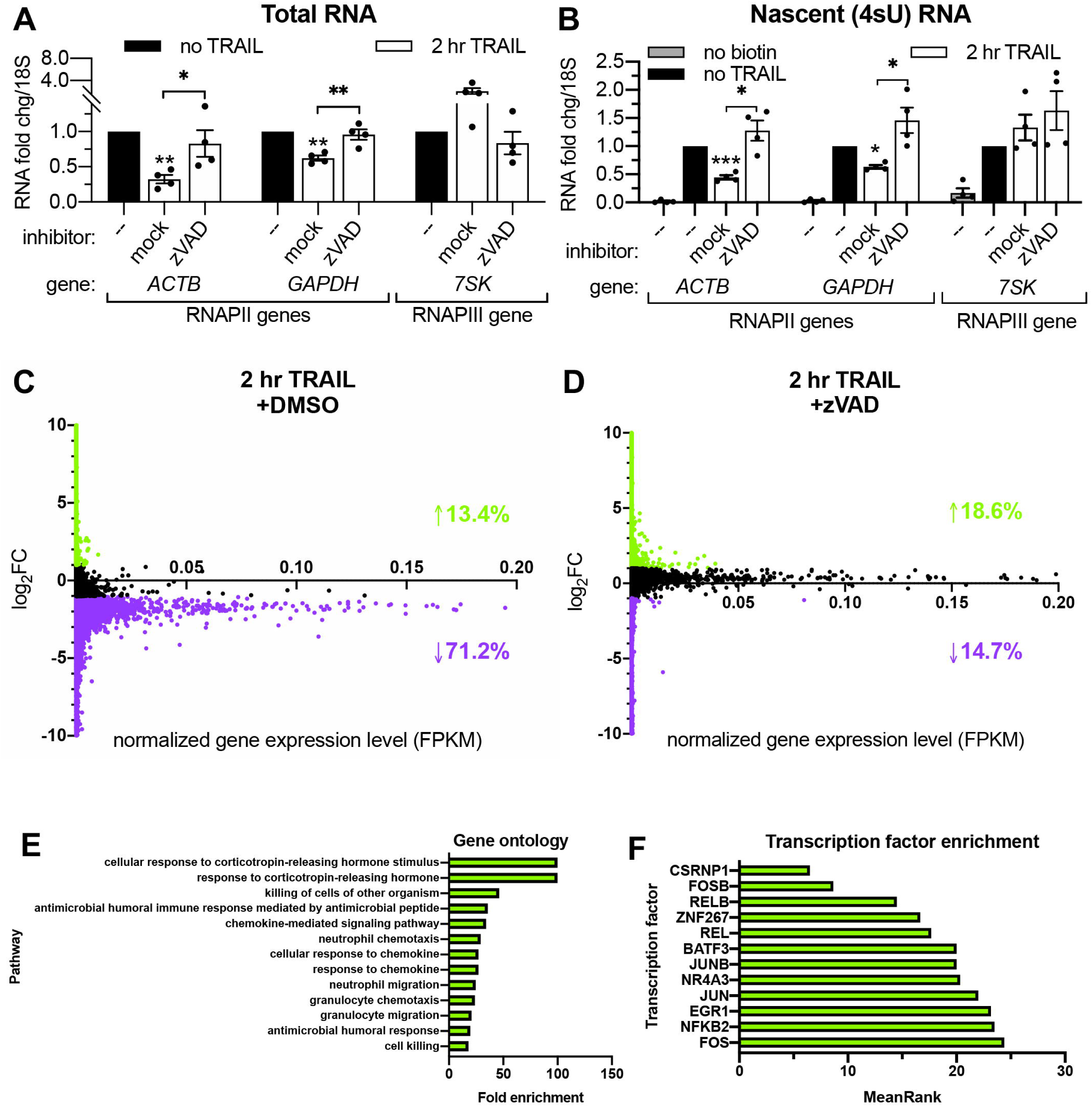
RNAPII transcription is globally repressed during early apoptosis. (A, B) RT-qPCR measurements of total (A) and nascent 4sU-labeled (B) RNA fold changes after 2 hr TRAIL treatment of HCT116 cells, including a 1 hr pre-treatment with either 40 μM zVAD or an equal volume of DMSO (“mock”). Also see Supplementary Figure 2A. RNA fold change values were calculated in reference to *18S* rRNA. Bar graphs display mean +/- SEM with individual biological replicates (*n* = 4) represented as dots. Statistically significant deviation from a null hypothesis of 1 was determined using one sample t test and indicated with asterisks directly above bars, while student’s t tests were performed to compare mean fold change values for mock inhibitor or scramble treated cells to those treated with inhibitor or a targeting siRNA and indicated with brackets. \**p*<0.05, \*\**p*<0.0.1, \*\*\**p*<0.001. (C, D) rRNA-depleted cDNA sequencing libraries were reverse transcribed from 4sU-labeled RNA isolated from cells under the conditions described in (A, B). Transcripts that aligned to genes in the human genome are graphed with differential log2 fold change expression values (log2FC) on the y axis and fragments per kilobase per million reads (FKPM) expression values on the x axis. All values were averaged from 2 biological replicates. Data points for transcripts upregulated or downregulated by 2-fold or greater are colored green and purple, respectively. Percentages of transcripts in each expression class are indicated with an arrow and in their corresponding colors. Also see Supplementary Figure 2C-D and Supplementary Tables 1-2. (E) Top statistically significant hits from gene ontology analysis performed for the list of (69) transcripts that were upregulated upon TRAIL treatment with DMSO in a statistically significant manner across biological duplicates. Also see Supplementary Table 3. (F) Top hits from transcription factor (TF) enrichment analysis for the same list of genes as above. The lower the MeanRank value, the more statistically significant enrichment for genes regulated by the indicated TF. Also see Supplementary Table 4.

Liu et al. (2018) reported that the mRNA degradation that occurs in HCT116 cells shortly after TRAIL treatment is widespread. In order to quantify the extent of the associated transcriptional arrest, we sequenced the 4sU-labeled nascent transcriptome of non-apoptotic and apoptotic cells. HCT116 cells were pre-treated with DMSO (i.e. allowing for apoptosis signaling to proceed) or zVAD (to prevent apoptosis) prior to addition of TRAIL (or vehicle) for 2 hr. Nascent RNA was depleted of rRNA before sequencing libraries were prepared, and >95% of the resultant reads mapped to mRNAs and other predominantly RNAPII transcripts such as lncRNAs (Marchese, Raimondi, & Huarte, 2017) and snoRNAs (Dieci, Preti, & Montanini, 2009) (Supplementary Tables 1 and 2). Significantly more nascent transcripts decreased than increased upon TRAIL treatment in cells pre-treated with DMSO (*p* ≅ 0, chi-squared test), while significantly more increased than decreased with TRAIL treatment in the presence of zVAD (*p* = 5.234e-143, chi-squared test). Markedly, 71.2% of the ∼28,000 unique transcripts detected were downregulated more than 2-fold in TRAIL-treated cells with DMSO (Figure 2C), while only 14.7% of ∼32,000 transcripts were repressed upon TRAIL treatment when caspases were inhibited with zVAD (Figure 2D). By contrast, fewer than 20% of transcripts were transcriptionally upregulated by the same amount during apoptosis in both conditions (Figure 2C-D). To validate that RNA production was accurately captured in the sequencing libraries, fold changes for a representative transcript (*ACTB*) in the original 4sU-labeled RNA samples were assessed by RT-qPCR using both exonic and intronic primers (Supplementary Figure 2B), showing good agreement between the three quantification methods.

Of the transcripts changed by 2-fold or greater in TRAIL-treated cells in the absence of zVAD, 745 decreased and 69 increased in a statistically significant manner across 2 biological replicates (Supplementary Figure 2C). By contrast, only 56 transcripts were significantly downregulated upon TRAIL treatment in the presence of zVAD, while 364 were upregulated (Supplementary Figure 2D). Gene ontology enrichment analysis revealed that the genes upregulated upon TRAIL treatment were disproportionately involved in cellular responses to chemokines and cell death (Figure 2E, Supplementary Table 3), while no statistically significant enrichments were observed in the downregulated transcripts (Supplementary Table 4). Interestingly, the induced genes were more likely to be regulated by transcription factors implicated in apoptosis such as CSRNP1 (Ye et al., 2017), FOSB (Baumann et al., 2003), NR4A3 (Fedorova et al., 2019), NFKB2 (Keller et al., 2010), JUNB (Gurzov, Ortis, Bakiri, Wagner, & Eizirik, 2008), JUN (Wisdom, Johnson, & Moore, 1999), EGR1 (Pignatelli, Luna-Medina, Pérez-Rendón, Santos, & Perez-Castillo, 2003), FOS (Preston et al., 1996), KFL6 (Huang, Li, & Guo, 2008), RELB (Guerin et al., 2002), and BATF3 (Qiu, Khairallah, Romanov, & Sheridan, 2020), indicating that the dataset reflects established apoptotic transcriptional dynamics (Figure 2F, Supplementary Table 5).

### Transcriptional repression during early apoptosis requires MOMP, but not necessarily caspase activity

We next sought to determine whether any hallmark features of apoptosis underlay the observed transcriptional repression. These include the limited proteolysis by the “initiator” CASP8, mass proteolysis by the “executioner” CASP3, the endonucleolytic cleavage of the genome by a caspase-activated DNase (CAD) that translocates into the nucleus during late stages of apoptosis (Enari et al., 1998) and MOMP (which instigates mRNA decay).

Small interfering RNA (siRNA) knockdowns (Figure 3A) were performed to first determine if the rescue of mRNA production caused by zVAD was due specifically to the inhibition of the initiator CASP8 or the executioner CASP3. Both accelerated mRNA decay (Supplementary Figure 3A) and RNAPII transcriptional repression (Figure 3B) upon 2 hr TRAIL treatment required the MOMP-inducing CASP8 but not CASP3, suggesting that mass proteolysis by CASP3 does not significantly contribute to either phenotype. Knockdown of CAD did not affect the reduction in 4sU incorporation observed during early apoptosis (Figure 3B), and DNA fragmentation (as measured by TUNEL assay) was not detected in TRAIL-treated HCT116 cells until 4-8 hr post-treatment (Supplementary Figure 3C). These findings are in agreement with the prior study showing that MOMP and mRNA decay occur before DNA fragmentation begins during extrinsic apoptosis (Thomas et al., 2015).

**Figure 3.**
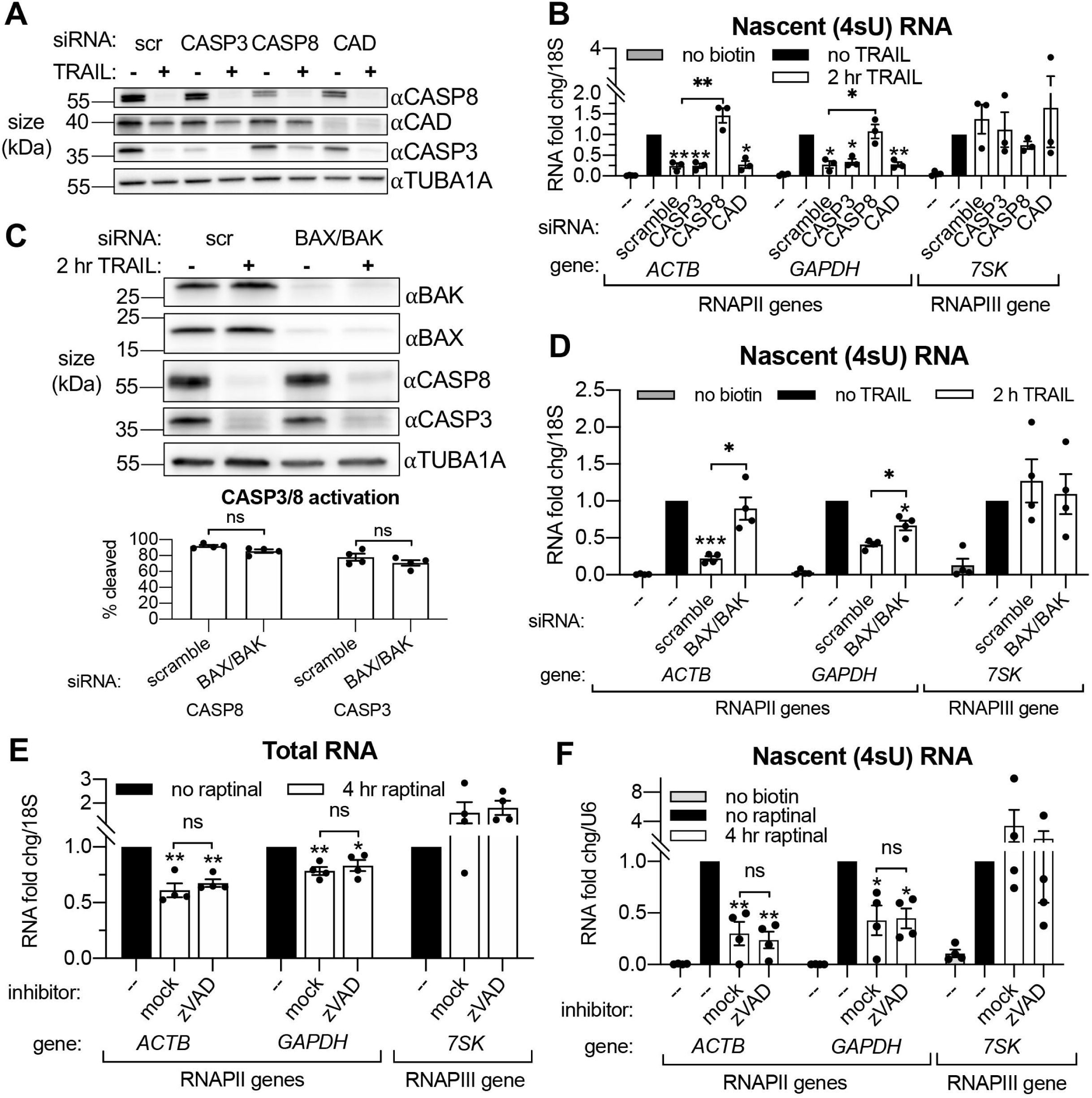
Transcriptional repression during early apoptosis requires MOMP, but not necessarily caspase activity. (A) Western blots showing the efficacy of CASP3, CASP8, and caspase-activated DNase (CAD) protein depletion following nucleofection with the indicated siRNAs, with or without 2 hr TRAIL treatment. α-Tubulin (TUBA1A) serves as a loading control. Blot representative of those from 4 biological replicates. (B) RT-qPCR measurements of 4sU-labeled nascent transcripts with or without 2 hr TRAIL treatment in cells nucleofected with the indicated the siRNAs (*n* = 3). Also see Supplementary Figures 3A-C. (C) Western blot detecting the indicated proteins in cells nucleofected with the indicated siRNA in the presence or absence of TRAIL. TUBA1A serves as a loading control. Blot representative of 4 biological replicates. The cleavage of CASP3 and CASP8 (as measured by the disappearance of the full-length form of each zymogen upon 2 hr TRAIL treatment, normalized to TUBA1A) is graphed below (*n* = 4). (D) 4sU-labeled RNA levels measured by RT-qPCR in HCT116 cells nucleofected with the indicated siRNAs, with or without 2 hr TRAIL treatment (*n* = 4). Also see Supplementary Figure 3D. (E, F) Total (E) and 4sU-labeled (F) RNA levels measured by RT-qPCR in HeLa cells after 10 μM raptinal treatment for 4 hr, with or without a 1 hr pre-treatment of 20 μM zVAD (*n* = 4). Also see Supplementary Figure 2E. Fold changes were calculated in reference to *U6* snRNA since its production was more stable after 4 hr raptinal than that of *18S* rRNA (see Supplementary Figure 2F). All RNA fold changes were calculated from Ct values normalized to *18S* or *U6* RNA, then normalized to non-apoptotic cells (“no TRAIL”) under otherwise identical conditions. Graphs display mean +/- SEM with individual biological replicates represented as dots. Statistically significant deviation from a null hypothesis of 1 was determined using one sample t test and indicated with asterisks directly above bars, while student’s t tests were performed to compare mean fold change values for mock inhibitor or scramble treated cells to those treated with zVAD or a targeting siRNA and indicated with brackets. The Holm-Sidak correction for multiple comparisons was applied in the student’s t tests represented in (B) \**p*<0.05, \*\**p*<0.0.1, \*\*\**p*<0.001.

Although CASP3 is dispensable for apoptotic transcriptional repression, a previous report suggests that CASP8 may cleave RPB1, the largest subunit of RNAPII (Lu, Luo, & Bregman, 2002). We therefore measured the protein expression of RPB1 and the 3 next largest RNAPII subunits (RBP2-4) during early apoptosis to determine if degradation of these subunits might explain the observed TRAIL-induced reduction in RNAPII transcription. Expression of RPB1-4 was relatively unaffected by 2 hr TRAIL treatment in the presence or absence of zVAD, with the exception of a small decrease in the amount of RPB2 that was rescued upon zVAD treatment (Supplementary Figure 3C). Thus, RNAPII depletion is unlikely to underlie the transcriptional repression phenotype.

Our above observations suggest that MOMP activation, for example through CASP8, is necessary to drive apoptotic mRNA decay and the ensuing transcriptional repression. To test this hypothesis, we attenuated MOMP in TRAIL-treated cells by depleting the mitochondrial pore-forming proteins BAX and BAK (Figure 3C). Indeed, siRNA-mediated depletion of BAX and BAK rescued cytoplasmic mRNA abundance (Supplementary Figure 3D) and RNAPII transcription (Figure 3D) of the *ACTB* and *GAPDH* transcripts in the presence of TRAIL, even though CASP8 and CASP3 were still cleaved (Figure 3D). Thus, CASP8 likely participates in this pathway only to the extent that it activates MOMP, as MOMP appears to be the main driver of mRNA decay and transcriptional repression.

Finally, to confirm that MOMP is sufficient to drive this phenotype, we used a small molecule, raptinal, that bypasses CASP8 to intrinsically induce MOMP (Heimer, Knoll, Schulze-Osthoff, & Ehrenschwender, 2019; Palchaudhuri et al., 2015). HeLa cells treated with 10 μM raptinal for 4 hr underwent MOMP, as measured by cytochrome c release into the cytoplasm, in the presence or absence of zVAD (Supplementary Figure 3E). Steady state levels (Figure 3E) and transcription (Figure 3F) of the aforementioned mRNAs was reduced upon raptinal treatment. The fact that this reduction in mRNA abundance and synthesis was maintained upon caspase inhibition by zVAD treatment indicates that the caspases are not required to drive these phenotypes outside of their role in MOMP activation.. Taken together, these data confirm that the mRNA degradation and ensuing transcriptional repression observed during early apoptosis are driven by MOMP.

### Cytoplasmic 3’- but not 5’-RNA exonucleases are required for apoptotic RNAPII transcriptional repression

Based on connections between virus-activated mRNA decay and RNAPII transcription (Abernathy et al., 2015; Gilbertson, Federspiel, Hartenian, Cristea, & Glaunsinger, 2018), we hypothesized that TRAIL-induced mRNA turnover was functionally linked to the concurrent transcriptional repression. Apoptotic mRNA decay occurs from the 3’ end by the actions of the cytoplasmic 3’-RNA exonuclease DIS3L2 and the mitochondrial 3’-RNA exonuclease PNPT1, which is released into the cytoplasm by MOMP (Liu et al., 2018; Thomas et al., 2015). This stands in contrast to basal mRNA decay, which occurs predominantly from the 5’ end by XRN1 (Jones, Zabolotskaya, & Newbury, 2012). We therefore set out determine if 3’ or 5’ decay factors were required for apoptosis-linked mRNA decay and the ensuing repression of mRNA transcription. Depletion of DIS3L2, PNPT1, or the cytoplasmic 3’ RNA exosome subunit EXOSC4 individually did not reproducibly rescue the total levels of either the *ACTB* or *GAPDH* mRNA during early apoptosis (Supplementary Figure 4A), nor did they affect the relative production of these transcripts (Supplementary Figure 4B). Given the likely redundant nature of the multiple 3’ end decay factors (Houseley & Tollervey, 2009), we instead performed concurrent knockdowns of DIS3L2, EXOSC4, and PNPT1 to more completely inhibit cytoplasmic 3’ RNA decay. We also knocked down the predominant 5’-3’ RNA exonuclease XRN1 to check the involvement of 5’ decay (Figure 4A). Depletion of the 3’-5’ but not the 5’-3’ decay machinery attenuated the apoptotic decrease in total RNA levels and largely restored RNAPII transcription (Figure 4B-C). Importantly, there was only a minor reduction in CASP3 activation in cells depleted of 3’-5’ decay factors and this was not significantly different from that observed upon XRN1 knockdown (Figure 4A). These observations suggest that decreased RNAPII transcription occurs as a consequence of accelerated 3’ mRNA degradation in the cytoplasm during early apoptosis.

**Figure 4.**
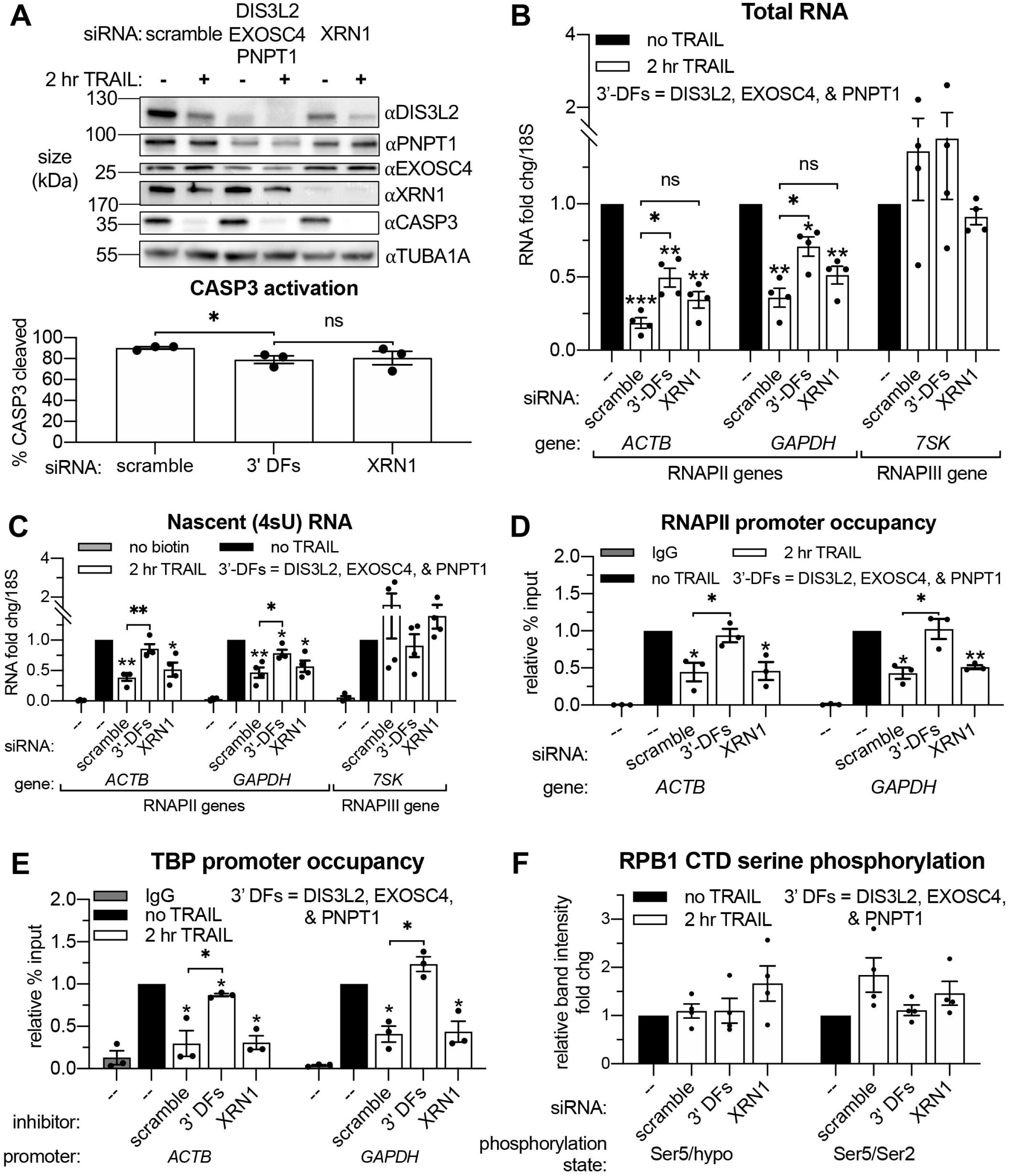
Apoptosis causes reduced RNAPII transcriptional output and promoter occupancy in an mRNA decay-dependent manner. (A) Western blots performed with HCT116 cell lysates depleted of the indicated decay factors with and without 2 hr TRAIL treatment. Blot representative of 3 biological replicates. Apoptosis induction was confirmed by disappearance of the full-length CASP3 band, quantified in the graph below by measuring band intensity normalized to an α-tubulin (TUBA1A) loading control (*n* = 3). (B, C) Changes in total (B) and nascent 4sU-labeled (*C*) RNA upon 2 hr TRAIL treatment in cells nucleofected with the indicated siRNAs were quantified by RT-qPCR (*n* = 4). Fold changes were calculated from Ct values normalized to *18S* rRNA. Also see Supplementary Figure 4A-B. (D, E) Chromatin immunoprecipitation (ChIP)-qPCR was used to measure occupancy of the indicated promoters by hypophosphorylated RNAPII (D) or TBP (E) under cellular conditions described in (A). Rabbit and mouse IgG antibodies were included in parallel immunoprecipitation reactions with chromatin from scramble siRNA-treated non-apoptotic cells in lieu of TBP and RNAPII antibodies, respectively, as a control. Also see Supplementary Figure 4D-E. (F) Relative band intensity ratios from 4 replicates of the representative western blots depicted in Supplementary Figure 4D, using primary antibodies specific to the indicated RPB1 CTD phosphorylation state under cellular conditions described in (A). Band intensity values were first normalized to a vinculin (VCL) loading control in each blot. All bar graphs display mean +/- SEM with individual biological replicates represented as dots. Statistically significant deviation from a null hypothesis of 1 was determined using one sample t test and indicated with asterisks directly above bars, while student’s t tests with the Holm-Sidak correction for multiple comparisons were performed to compare mean values between groups indicated with brackets. \**p*<0.05, \*\**p*<0.0.1, \*\*\**p*<0.001.

### Apoptosis causes reduced RNAPII promoter occupancy in an mRNA decay-dependent manner

RNAPII is recruited to promoters in an unphosphorylated state, but its subsequent promoter escape and elongation are governed by a series of phosphorylation events in the heptad (Y_1_S_2_P_3_T_4_S_5_P_6_S_7_)_n_ repeats of the RPB1 C-terminal domain (CTD). To determine the stage of transcription impacted by mRNA decay, we performed RNAPII ChIP-qPCR and western blots using antibodies recognizing the different RNAPII phosphorylation states. Occupancy of hypophosphorylated RNAPII at the *ACTB* and *GAPDH* promoters was significantly reduced after 2 hr TRAIL treatment (Figure 3D). In accordance with the 4sU labeling results, siRNA-mediated knockdowns showed that loss of RNAPII occupancy in response to TRAIL requires CASP8 but not CASP3 (Supplementary Figure 4C), as well as cytoplasmic 3’-5’ RNA decay factors but not the 5’-3’ exonuclease XRN1 (Figure 4D). Impaired binding of the TATA-binding protein (TBP), which nucleates the formation of the RNAPII pre-initiation complex (PIC) at promoters (Buratowski, Hahn, Guarente, & Sharp, 1989; Louder et al., 2016), mirrored that of RNAPII (Figure 4E). These changes are not driven by alterations in the stability of RPB1 or TBP, since the expression of these proteins remained relatively constant during early apoptosis regardless of the presence of mRNA decay factors (Supplementary Figure 4D-E). The relative proportion of initiating RPB1 CTD phosphorylated at the serine 5 position and the ratio of serine 5 to serine 2 phosphorylation, which decreases during transcriptional elongation (Shandilya & Roberts, 2012), also remain unchanged during early apoptosis regardless of the presence of mRNA decay factors (Figure 4F). These data suggest that the decay-dependent reduction in mRNA synthesis occurs at or before PIC formation, rather than during transcriptional initiation and elongation.

### Importin α/β transport links mRNA decay and transcription

Finally, we sought to determine how TRAIL-induced cytoplasmic mRNA decay signals to the nucleus to induce transcriptional repression. Data from viral systems suggest that this signaling involves differential trafficking of RNA binding proteins (RBPs), many of which transit to the nucleus in response to virus-induced cytoplasmic mRNA decay (Gilbertson et al., 2018). We therefore sought to test the hypothesis that nuclear import of RBPs underlies how cytoplasmic mRNA decay is sensed by the RNAPII transcriptional machinery.

To determine whether RBP redistribution occurs during early apoptosis, we analyzed the subcellular distribution of cytoplasmic poly(A) binding protein PABPC1, an RBP known to shuttle to the nucleus in response to virus-induced RNA decay (Gilbertson et al., 2018; Kumar & Glaunsinger, 2010; Lee & Glaunsinger, 2009). We performed cell fractionations of HCT116 cells to measure PABPC1 levels in the nucleus versus the cytoplasm upon induction of apoptosis. Indeed, 2 hr after TRAIL treatment, PABPC1 protein levels increased in the nuclear fraction of cells but not in the cytoplasmic fraction, indicative of relocalization (Figure 5A). This increase in nuclear PABPC was dependent on the presence of CASP8 but not CASP3, mirroring the incidence of mRNA decay and reduced RNAPII transcription. Thus, similar to viral infection, PABPC1 relocalization occurs coincidentally with increased mRNA decay and transcriptional repression in the context of early apoptosis.

**Figure 5.**
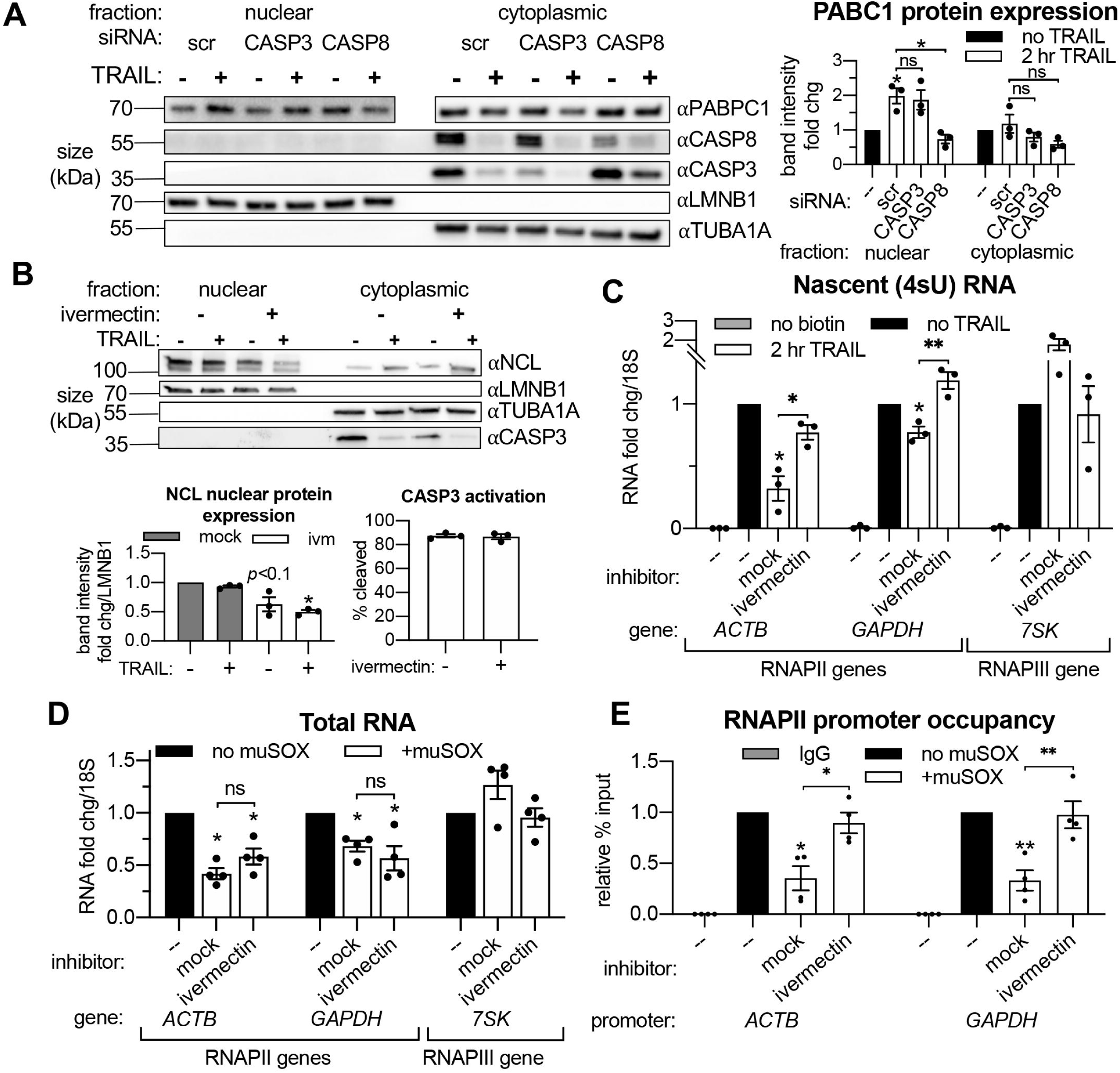
Importin α/β transport links mRNA decay and transcription. (A) Western blots showing the indicated proteins in the nuclear and cytoplasmic fractions of apoptotic and non-apoptotic HCT116 cells nucleofected with the indicated siRNAs. The nuclear and cytoplasmic fractions of PABPC1 were imaged on the same membrane section but cropped and edited separately to visualize the low nuclear expression of this canonically cytoplasmic protein. Protein expression (right) was calculated by band intensity in reference to lamin B1 (LMNB1) or α-tubulin (TUBA1A) loading controls, for the nuclear and cytoplasmic fractions, respectively. (B) Nuclear and cytoplasmic expression of the indicated proteins in apoptotic and non-apoptotic cells with a 1 hr pre-treatment with 25 M ivermectin or an equal volume of EtOH (“mock”). Nuclear levels of the importin α/β substrate nucleolin (NCL) were quantified by band intensity in reference to a LMNB1 loading control, while disappearance of full-length CASP3 was quantified in the cytoplasm in reference to TUBA1A. Also see Supplementary Figure 5A (C) Levels of nascent 4sU-labeled RNA were quantified by RT-qPCR under the cellular conditions described in (B). Also see Supplementary Figure 5B-C. (D) RT-qPCR quantification of total RNA levels in HEK293T cells stably expressing a doxycycline (dox)-inducible form of muSOX endonuclease, cultured with or without 1 μg/mL dox for 24 hr. Cells were treated with 25 μM ivermectin or an equal volume of EtOH 2 hr before harvesting. (E) RNAPII promoter occupancy at the ACTB and GAPDH promoters was determined by ChIP-qPCR under cellular conditions described in (D). Also see Supplementary Figure 5D-F. RNA fold changes were calculated from Ct values normalized to 18S rRNA. All bar graphs display mean +/- SEM with individual biological replicates represented as dots. Statistically significant deviation from a null hypothesis of 1 was determined using one sample t test and indicated with asterisks directly above bars, while student’s t tests were performed to compare mean fold change values for mock inhibitor or scramble treated cells to those treated with ivermectin or a targeting siRNA and indicated with brackets. The Holm-Sidak correction for multiple comparisons was applied in the student’s t tests represented in (A). *p<0.05, **p<0.0.1, ***p<0.001.

We next sought to directly evaluate the role of protein shuttling in connecting cytoplasmic mRNA decay to transcriptional repression. The majority of proteins ∼60 kilodaltons (kDa) and larger cannot passively diffuse through nuclear pores; they require assistance by importins to enter the nucleus from the cytoplasm (Gö, 1998). Classical nuclear transport occurs by importin α binding to a cytoplasmic substrate, which is then bound by an importin β to form a tertiary complex that is able to move through nuclear pore complexes (Stewart, 2007). To test if this mode of nuclear transport is required for transcriptional feedback, HCT116 cells were pretreated with ivermectin, a specific inhibitor of importin α/β transport (Wagstaff, Sivakumaran, Heaton, Harrich, & Jans, 2012), before the 2 hr TRAIL treatment. The efficacy of ivermectin was validated by an observed decrease in the nuclear levels of the RBP nucleolin (NCL), a known importin α/β substrate (Figure 5B). Ivermectin pretreatment rescued RNAPII transcription upon TRAIL treatment (Figure 5C) without significantly affecting the extent of mRNA decay (Supplementary Figure 5A), suggesting that protein trafficking to the nucleus provides signals connecting cytoplasmic mRNA decay to transcription.

Importantly, ivermectin did not diminish the extent of CASP3 cleavage during early apoptosis (Figure 5B), nor did it decrease baseline levels of transcription in non-apoptotic cells (Supplementary Figure 5B). Interestingly, it also did not prevent PABPC1 import (Supplementary Figure 5C), suggesting that PABPC1 translocation likely occurs via an importin α/β-independent pathway and is not sufficient to repress RNAPII transcription.

Finally, we evaluated whether importin α/β transport is also required for feedback between viral nuclease-driven mRNA decay and RNAPII transcription, as this would suggest that the underlying mechanisms involved in activating this pathway may be conserved. We used the mRNA specific endonuclease muSOX from the gammaherpesvirus MHV68, as muSOX expression has been shown to cause widespread mRNA decay and subsequent transcriptional repression (Abernathy et al., 2014, 2015). HEK-293T cell lines were engineered to stably express dox-inducible wild-type muSOX or the catalytically inactive D219A mutant (Abernathy et al., 2015). These cells were treated with ivermectin for 3 hr and the resultant changes in RNAPII promoter occupancy were measured by ChIP-qPCR. As expected, expression of WT (Figure 5D-E) but not D219A muSOX (Supplementary Figure 5E-F) caused mRNA decay and transcriptional repression. Notably, inhibiting nuclear import with ivermectin (Supplementary Figure 5D) rescued RNAPII promoter occupancy (Figure 5E) without altering the extent of mRNA decay in muSOX expressing cells (Figure 5D). Thus, importin α/β nuclear transport plays a key role in linking cytoplasmic mRNA decay to nuclear transcription, both during early apoptosis and upon viral nuclease expression.

## DISCUSSION

mRNA decay and synthesis rates are tightly regulated in order to maintain appropriate levels of cellular mRNA transcripts (Braun & Young, 2014). It is well established that when cytoplasmic mRNA is stabilized, for example by the depletion of RNA exonucleases, transcription often slows in order to compensate for increased transcript abundance (Haimovich et al., 2013; Helenius et al., 2011; Singh et al., 2019; Sun et al., 2012). Eukaryotic cells thus have the capacity to “buffer” against reductions in mRNA turnover or synthesis. Here, we revealed that a buffering response does not occur under conditions of elevated cytoplasmic mRNA degradation stimulated during early apoptosis. Instead, cells respond to cytoplasmic mRNA depletion by decreasing RNAPII promoter occupancy and transcript synthesis, thereby amplifying the magnitude of the gene expression shut down. Nuclear import of cytoplasmic proteins is required for this “transcriptional feedback”, suggesting a pathway of gene regulation in which enhanced mRNA decay prompts cytoplasmic proteins to enter the nucleus and halt mRNA production. Notably, similar transcriptional feedback is elicited during virus-induced mRNA decay (Abernathy et al., 2015; Gilbertson et al., 2018; Hartenian et al., 2020), indicating that distinct cellular stresses can converge on this pathway to potentiate a multi-tiered shutdown of gene expression.

Multiple experiments support the conclusion that the TRAIL-induced transcriptional repression phenotype is a consequence cytoplasmic decay triggered by MOMP, rather than caspase activity. XRN1-driven 5’-3’ end decay is the major pathway involved in basal mRNA decay (Łabno, Tomecki, & Dziembowski, 2016), but MOMP-induced mRNA decay is primarily driven by 3’ exonucleases such as PNPT1 and DIS3L2 (Liu et al., 2018; Thomas et al., 2015). Accordingly, co-depletion of 3’ decay factors but not XRN1 restored RNAPII promoter occupancy and transcription during early apoptosis. In contrast to that of 3’ mRNA decay factors, depletion of CASP3 (or the caspase activated DNase CAD) did not block mRNA degradation or transcriptional repression, even though CASP3 is responsible for the vast majority of proteolysis that is characteristic of apoptotic cell death (J. G. Walsh et al., 2008). Additionally, the initiator CASP8 was required only under conditions where its activity was needed to induce MOMP. These findings reinforce the idea that mRNA decay and transcriptional repression are very early events that are independent of the subsequent cascade of caspase-driven phenotypes underlying many of the hallmark features of apoptosis.

A key open question is what signal conveys cytoplasmic mRNA degradation information to the nucleus to cause transcriptional repression. Our data are consistent with a model in which the signal is provided by one or more proteins imported into the nucleus in response to accelerated mRNA decay (Figure 6). Indeed, many cytoplasmic RNA binding proteins undergo nuclear-cytoplasmic redistribution under conditions of viral nuclease-induced mRNA decay, including PABPC (Gilbertson et al., 2018; Kumar & Glaunsinger, 2010; Kumar, Shum, & Glaunsinger, 2011). We propose that a certain threshold of mRNA degradation is necessary to elicit protein trafficking and transcriptional repression. Presumably, normal levels of basal mRNA decay and regular cytoplasmic repopulation result in a balanced level of mRNA-bound versus unbound proteins. However, if this balance is tipped during accelerated mRNA decay, an excess of unbound RNA binding proteins could accumulate and be transported into the nucleus.

**Figure 6.**
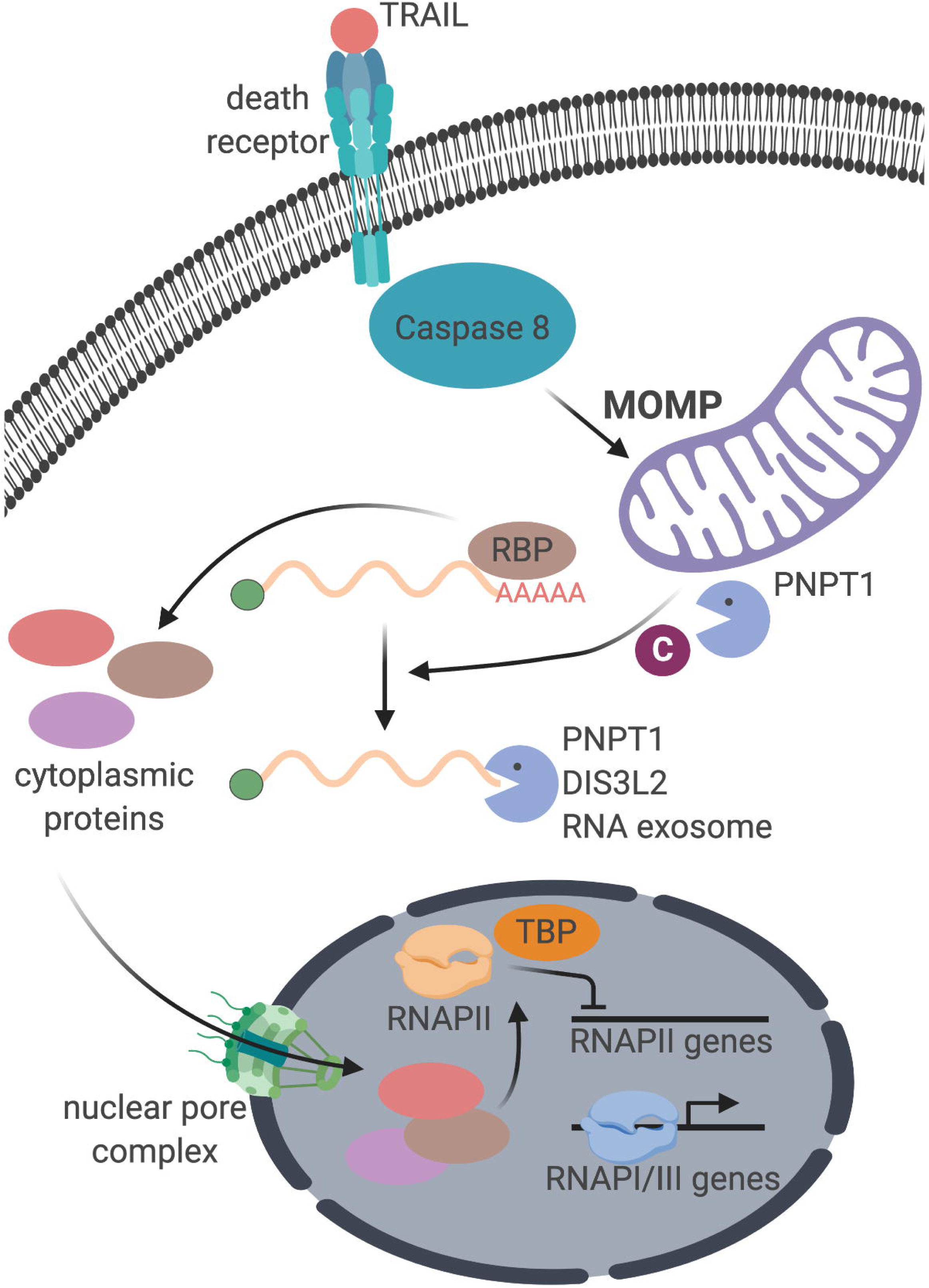
Schematic representation of the cellular events connecting apoptosis induction with mRNA decay and RNAPII transcription.

PABPC1 is likely among the first proteins released from mRNA transcripts undergoing 3’ end degradation, and its nuclear translocation is driven by a poly(A)- masked nuclear localization signal that is exposed upon RNA decay (Kumar et al., 2011). Indeed, we found that PABPC1 undergoes nuclear translocation during early apoptosis. However, the fact that ivermectin blocks transcriptional repression but not PABPC1 import indicates that while PABPC1 redistribution is a marker of mRNA decay, it is not sufficient to induce transcriptional repression in this system. Instead, we hypothesize that the observed RNAPII transcriptional repression occurs as a result of the cumulative nuclear import of multiple factors via importin α/β. Our group previously reported that at least 66 proteins in addition to PABPC1 are selectively enriched in the nucleus upon transfection with the viral endonuclease muSOX (but not with the catalytically-dead D219A mutant), 22 of which are known to be RNA-binding proteins and 45 of which shuttle in a manner dependent on the cytoplasmic mRNA exonuclease primarily responsible for clearing endonuclease cleavage fragments, XRN1 (Gilbertson et al., 2018). Future studies in which changes in the nuclear and cytoplasmic proteome upon apoptosis induction and viral endonuclease expression in the presence and absence of ivermectin will likely provide insight into which additional protein or proteins play a role in connecting cytoplasmic mRNA turnover to RNAPII transcription. Nonetheless, the requirement for importin α/β−mediated nuclear transport in both apoptosis-induced and viral nuclease-induced transcriptional repression suggests that these two stimuli may activate a conserved pathway of gene regulation. Whether other types of cell stress elicit a similar response remains an important question for future investigation.

The mRNA 3’ end decay-dependent decrease in RNA synthesis is accompanied by a reduction in TBP and RNAPII occupancy at promoters, consistent with transcriptional inhibition occurring upstream of RNAPII initiation and elongation (Gilbertson et al., 2018; Hartenian et al., 2020). The mechanism driving such a defect is yet to be defined, but possibilities include changes to transcript processing pathways, transcriptional regulators, or the chromatin state that directly or indirectly impact formation of the preinitiation complex. In this regard, mRNA processing and transcription are functionally linked (Bentley, 2014) and nuclear accumulation of PABPC1 has been shown to affect mRNA processing by inducing hyperadenylation of nascent transcripts (Kumar & Glaunsinger, 2010; Lee & Glaunsinger, 2009). Furthermore, TBP interacts with the cleavage–polyadenylation specificity factor (Dantonel, Murthy, Manjey, & Tora, 1997), providing a possible link between RNAPII preinitiation complex assembly and polyadenylation. Such a mechanism could also explain the specificity of the observed transcriptional repression to RNAPII transcripts. Chromatin architecture could also be influenced by shuttling of RNA binding proteins, as for example the yeast nucleocytoplasmic protein Mrn1 is implicated in the function of chromatin remodeling complexes (Düring et al., 2012). Interestingly, the polycomb repressive chromatin complex 2 (PRC2) and DNA methyltransferase 1 (DNMT1) have been shown to bind both nuclear mRNA and chromatin in a mutually exclusive manner, (Beltran et al., 2016; Di Ruscio et al., 2013; Garland et al., 2019), evoking the possibility that a nuclear influx of RBPs could secondarily increase the chromatin association of such transcriptional repressors. Future work will test these possibilities in order to mechanistically define the connection between nuclear import and RNAPII transcription under conditions of enhanced mRNA decay.

There are several potential benefits to dampening mRNA transcription in response to accelerated mRNA turnover. Debris from apoptotic cells is usually cleared by macrophages, but inefficient clearance of dead cells can lead to the development of autoantibodies to intracellular components such as histones, DNA, and ribonucleoprotein complexes. This contributes to autoimmune conditions such as systemic lupus erythematosus (Caruso & Poon, 2018; Nagata, Hanayama, & Kawane, 2010). In the context of infection, many viruses require the host RNAPII transcriptional machinery to express viral genes (Harwig et al., 2017; Rivas, Schmaling, & Gaglia, 2016; Walker & Fodor, 2019). It may therefore be advantageous for the cell to halt transcription in attempt to pre-empt a viral takeover of mRNA synthesis. That said, as with antiviral translational shutdown mechanisms (D. Walsh et al., 2013), viruses have evolved strategies to evade transcriptional repression (Hartenian et al., 2020; Harwig et al., 2017), as well as inhibit cell death via apoptosis and/or co-opt apoptotic signaling cascades (Suffert et al., 2011; Tabtieng, Degterev, & Gaglia, 2018; Zhang, Hildreth, & Colberg-Poley, 2013; Zhou, Jiang, Liu, Liu, & Liang, 2017). In either case, the shutdown of transcription in a cell under stress may protect surrounding cells from danger, particularly in an in vivo context. If true, this type of response is likely to be more relevant in multicellular compared with unicellular organisms.

## ACKNOWLEDGEMENTS

This work was supported by NIH grant R01CA136367 to B.G. and the HHMI Gilliam Fellowship for Advanced Study to C.D. B.G. is an investigator of the Howard Hughes Medical Institute. We thank the UC Berkeley Cell Culture Facility for providing the cell lines used in this study, in addition to the UC Berkeley DNA Sequencing Facility and all members of the Glaunsinger and Coscoy labs for providing valuable feedback.

## AUTHOR CONTRIBUTIONS

C.D. conceived and performed the experiments, analyzed and visualized the data, acquired funding, and drafted the manuscript. E.H. created the stable cell lines and drafted the corresponding section of the manuscript, as well as directed and assisted with analysis of the 4sU-seq data. V.K. analyzed the 4sU-seq data. B.G. conceived and supervised the study, acquired funding, and edited the manuscript.

## DECLARATION OF INTERESTS

The authors declare no competing interests.

## FIGURE LEGENDS

**Supplementary Figure 1.**
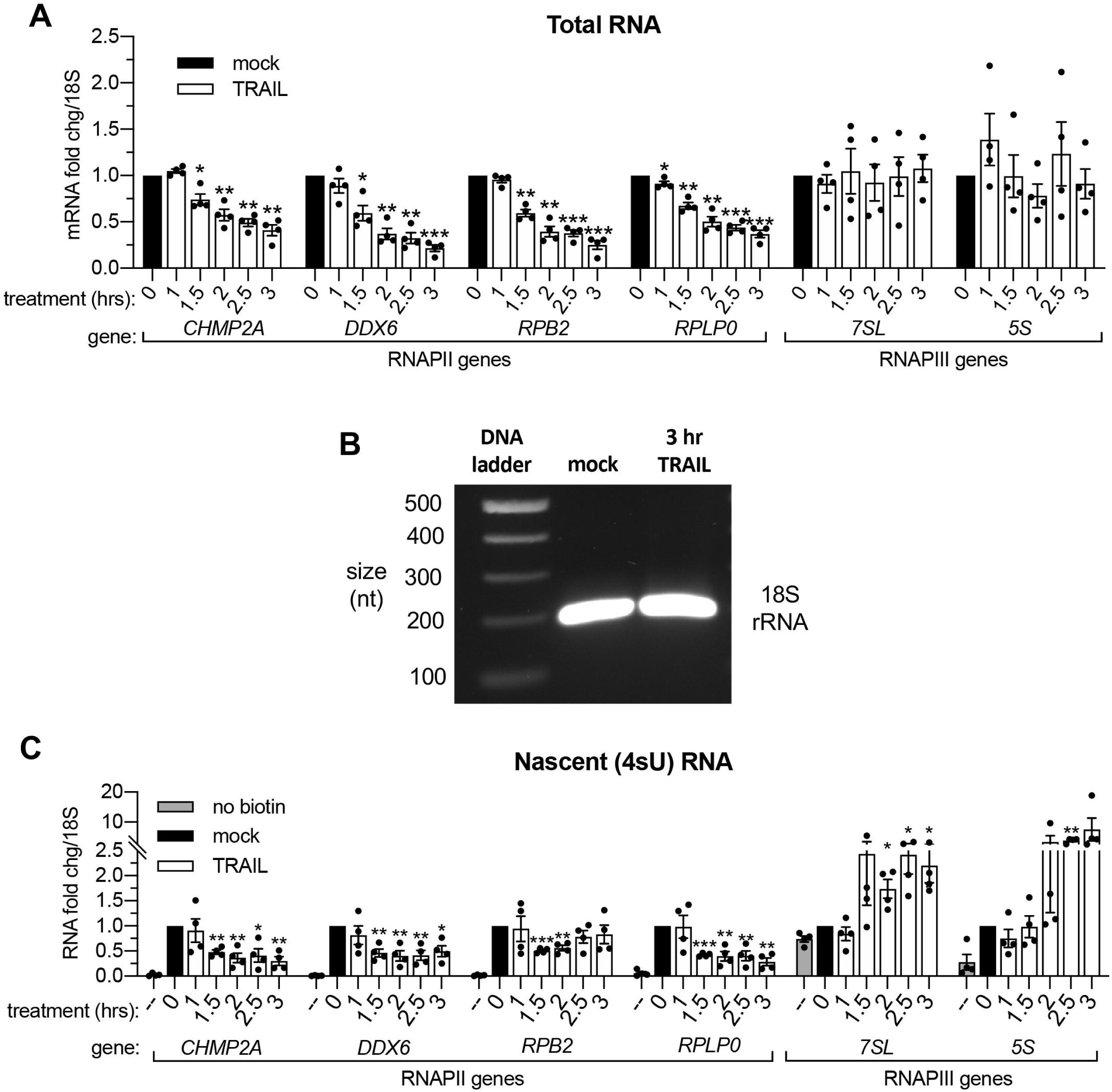
mRNA decay during early apoptosis is accompanied by reduced synthesis of RNAPII transcripts. (A) RT-qPCR quantification of total RNA at the indicated times post 100 ng/μL TRAIL treatment of HCT116 cells (*n* = 4). (B) Stained agarose gel depicting a 200 nt RT-PCR product of 4sU-labeled 18S rRNA, extracted and isolated from an equal number of cells treated with 3 hr vehicle (“mock”) or 100 ng/μL TRAIL. Gel representative of that from 3 biological replicates. (C) RT-qPCR quantification of 4sU-labeled RNA at the indicated times post 100 ng/μL TRAIL treatment of HCT116 cells (*n* = 4). Fold changes were calculated from Ct values normalized to *18S* rRNA in reference to mock treated cells. Graphs display mean +/- SEM with individual biological replicates represented as dots. Statistically significant deviation from a null hypothesis of 1 was determined using one sample t test; \**p*<0.05, \*\**p*<0.01, \*\*\**p*<0.001.

**Supplementary Figure 2.**
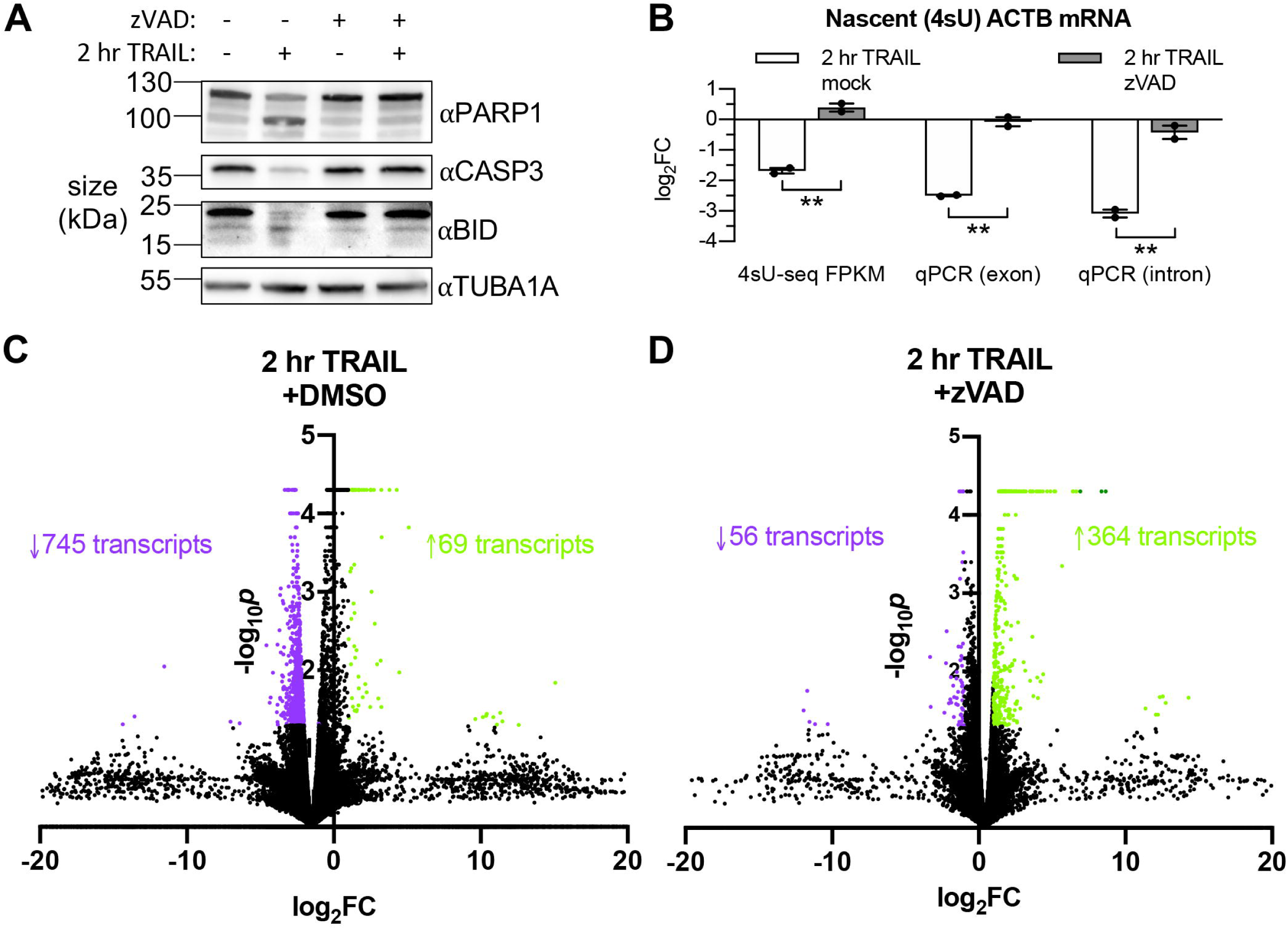
RNAPII transcription is globally repressed during early apoptosis. (A) Western blot of HCT116 lysates showing cleavage (or lack thereof) of the caspase 8 (CASP8) targets BID and caspase 3 (CASP3), as well as the CASP3 target PARP1 upon 2 hr 100 ng/μL TRAIL treatment in the presence or absence of 40 μM zVAD. Blot representative of that from 3 biological replicates. (B) Log2 fold change (log_2_FC) in abundance of the ACTB transcript upon 2 hr TRAIL treatment in the presence of either zVAD or DMSO vehicle (“mock”). Fold changes from the same 4sU- labeled RNA samples were quantified by next-generation sequencing and differential expression analysis, as well as RT-qPCR normalized to 18S rRNA using both intronic and exonic gene-specific primers. Graph displays mean +/- SEM with individual biological replicates (*n* = 2) represented as dots. Student’s t tests were performed to compare the log_2_FC values upon TRAIL treatment in the presence of DMSO vehicle or zVAD. \*\**p*<0.0.1 (C, D) Volcano plot depicting the –log_10_*p* values across biological duplicates for the fragment per kilobase per million read fold changes in the sequencing libraries depicted in Figures 2C, D. Each point represents a transcript mapped to the human genome. Upregulated and downregulated transcripts with *p* < 0.05 are colored in green and purple, respectively. Number of transcripts in each expression class are indicated with an arrow and in their corresponding colors.

**Supplementary Figure 3.**
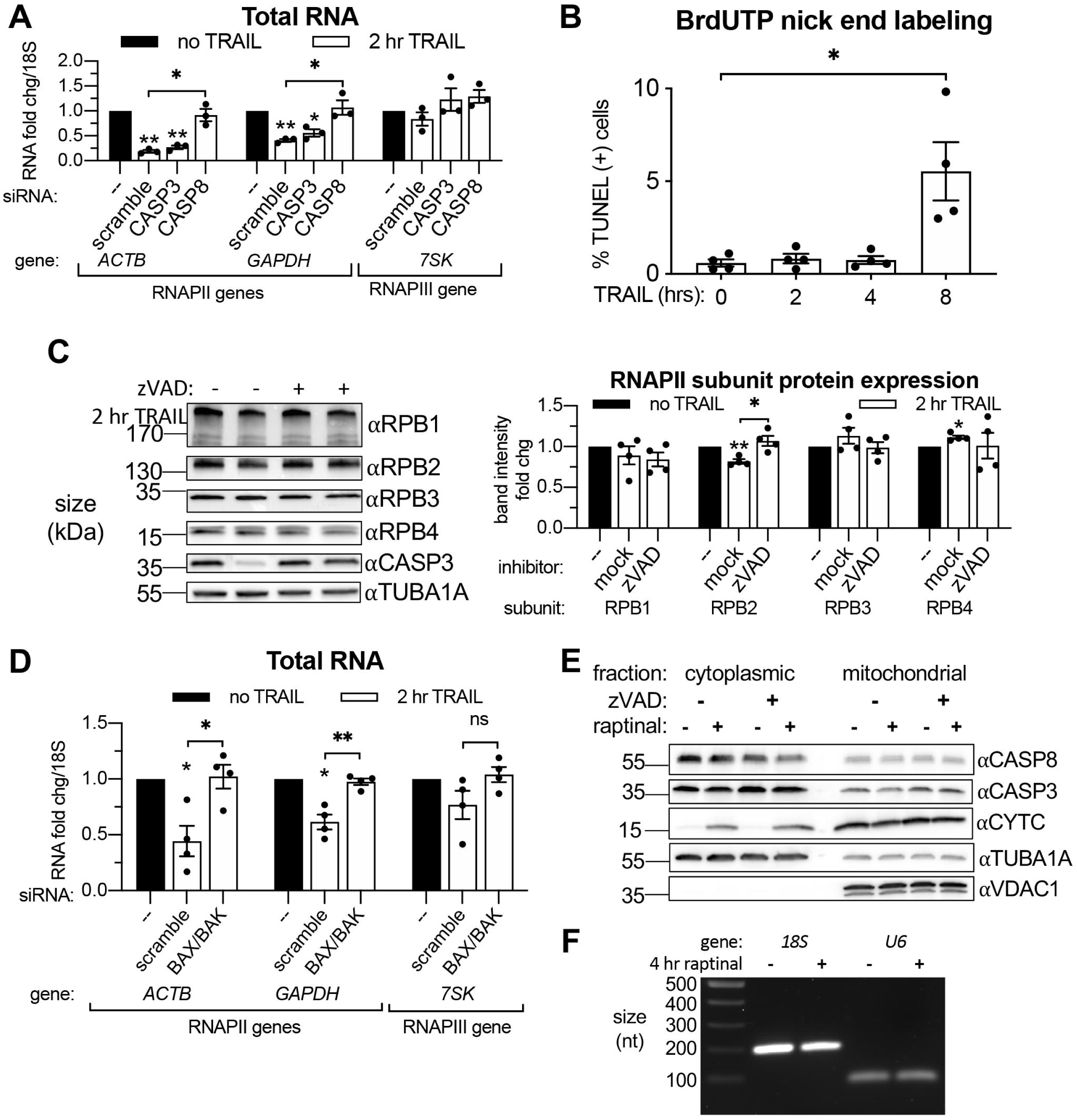
Transcriptional repression during early apoptosis requires MOMP, but not necessarily caspase activity. (A) RT-qPCR quantification of the change in total RNA levels in HCT116 cells treated with the indicated siRNAs upon 2 hr TRAIL treatment. Fold change were calculated from Ct values normalized to *18S* rRNA. (B) Quantification of BrdUTP-labeled HCT116 cells treated with TRAIL for the indicated times in a TUNEL assay (*n* = 4). Percent positive cells calculated by flow cytometry using FloJo imaging software to count the number of cells with elevated FL2 fluorescence. (C) Western blot of HCT116 lysates from cells under conditions described in (A) depicting levels of the indicated RNAPII subunits and CASP3 activation. α-Tubulin (TUBA1A) serves as a loading control. Graph to the right quantifies the change in RNAPII subunit levels upon apoptosis induction by TUBA1A-normalized band density. (D) Total RNA levels measured by RT-qPCR in HCT116 cells nucleofected with the indicated siRNAs, with or without 2 hr TRAIL treatment (*n* = 4). (E) Western blot of the cytoplasmic and mitochondrial fractions of HeLa cells treated with 10 μM raptinal or equal volume of DMSO, in the presence of 20 μM zVAD or additional volume of DMSO. Efficacy of raptinal treatment was demonstrated by cytochrome c (CYTC) release into the cytoplasm in the presence and absence of zVAD, which effectively blocked the cleavage and activation of CASP3. TUBA1A and VDAC1 served as cytoplasmic and mitochondrial loading controls, respectively. Blot representative of that from 3 biological replicates. (F) Stained agarose gel depicting a 200 nt and 101 nt RT-PCR product of 4sU-labeled *18S* and *U6* rRNA, respectively, extracted and isolated from an equal number of cells treated with 4 hr 10 μM raptinal or equal volume of DMSO. Gel representative of that from 3 biological replicates. All bar graphs display mean +/- SEM with individual biological replicates represented as dots. Statistically significant deviation from a null hypothesis of 1 was determined using one sample t test and indicated with asterisks directly above bars, while student’s t tests were performed to compare mean fold change values for mock inhibitor or scramble treated cells to those treated with zVAD or a targeting siRNA and indicated with brackets. The Holm-Sidak correction for multiple comparisons was applied in the student’s t tests represented in (B). \**p*<0.05, \*\**p*<0.0.1, \*\*\**p*<0.001.

**Supplementary Figure 4.**
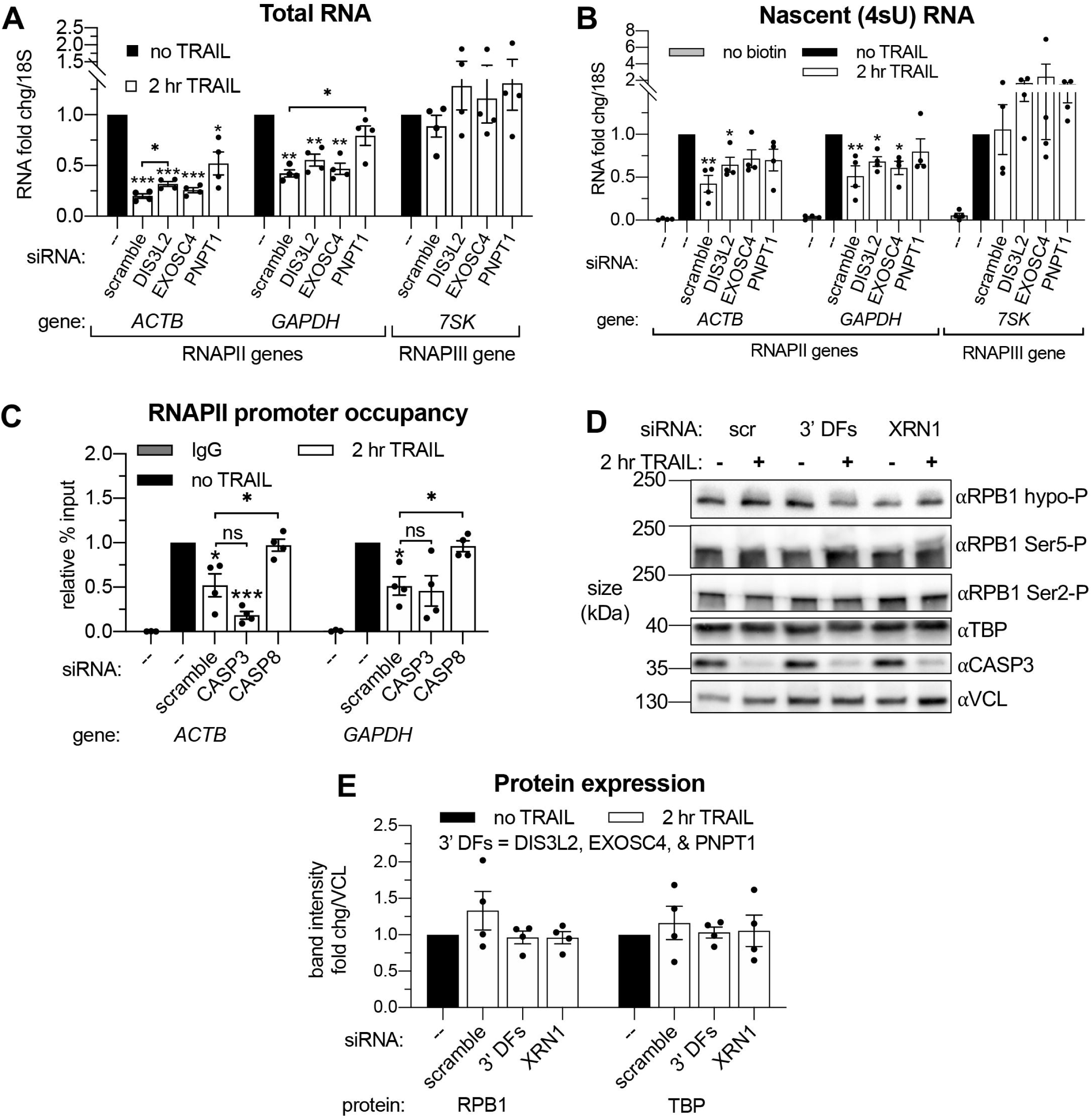
Apoptosis causes reduced RNAPII transcriptional output and promoter occupancy in an mRNA decay-dependent manner. (A, B) RT-qPCR quantification of the change in total (A) and 4sU-labeled (B) RNA levels in HCT116 cells treated with the indicated siRNAs upon 2 hr TRAIL treatment (*n* = 4). Fold changes were calculated from Ct values normalized to *18S* rRNA. (C) Chromatin immunoprecipitation (ChIP)-qPCR was used to measure the change in hypophosorylated RNAPII occupancy at the indicated promoters in cells nucleofected with the indicated siRNAs (*n* = 4). Mouse IgG antibody was included in parallel immunoprecipitation reactions with chromatin from scramble siRNA-treated non-apoptotic cells in lieu of RNAPII antibody as a control. (D) Representative western blots on lysates of HCT116 cells depleted of either 3’ DFs (DIS3L2, EXOSC4, and PNPT1) or XRN1 before and after 2 hr TRAIL treatment, probing for the indicated protein or RPB1 C-terminal domain (CTD) phosphorylation state. Apoptosis induction was confirmed by observing CASP3 cleavage. A vinculin (VCL) loading control was imaged for the blots of each phosphorylation state, but a single representative panel is shown. Quantification of RPB1 phosphorylation states from 4 replicates displayed in Figure 4F. (E) Band intensity fold changes of hypophosphorylated RPB1 and TBP derived from 4 biological replicates of the experiment described in (D). Bar graph displays mean +/- SEM with individual biological replicates represented as dots. Statistically significant deviation from a null hypothesis of 1 was determined using one sample t test and indicated with asterisks directly above bars, while student’s t tests with the Holm-Sidak correction for multiple comparisons were performed to compare mean values between groups indicated with brackets. \**p*<0.05, \*\**p*<0.0.1, \*\*\**p*<0.001.

**Supplementary Figure 5.**
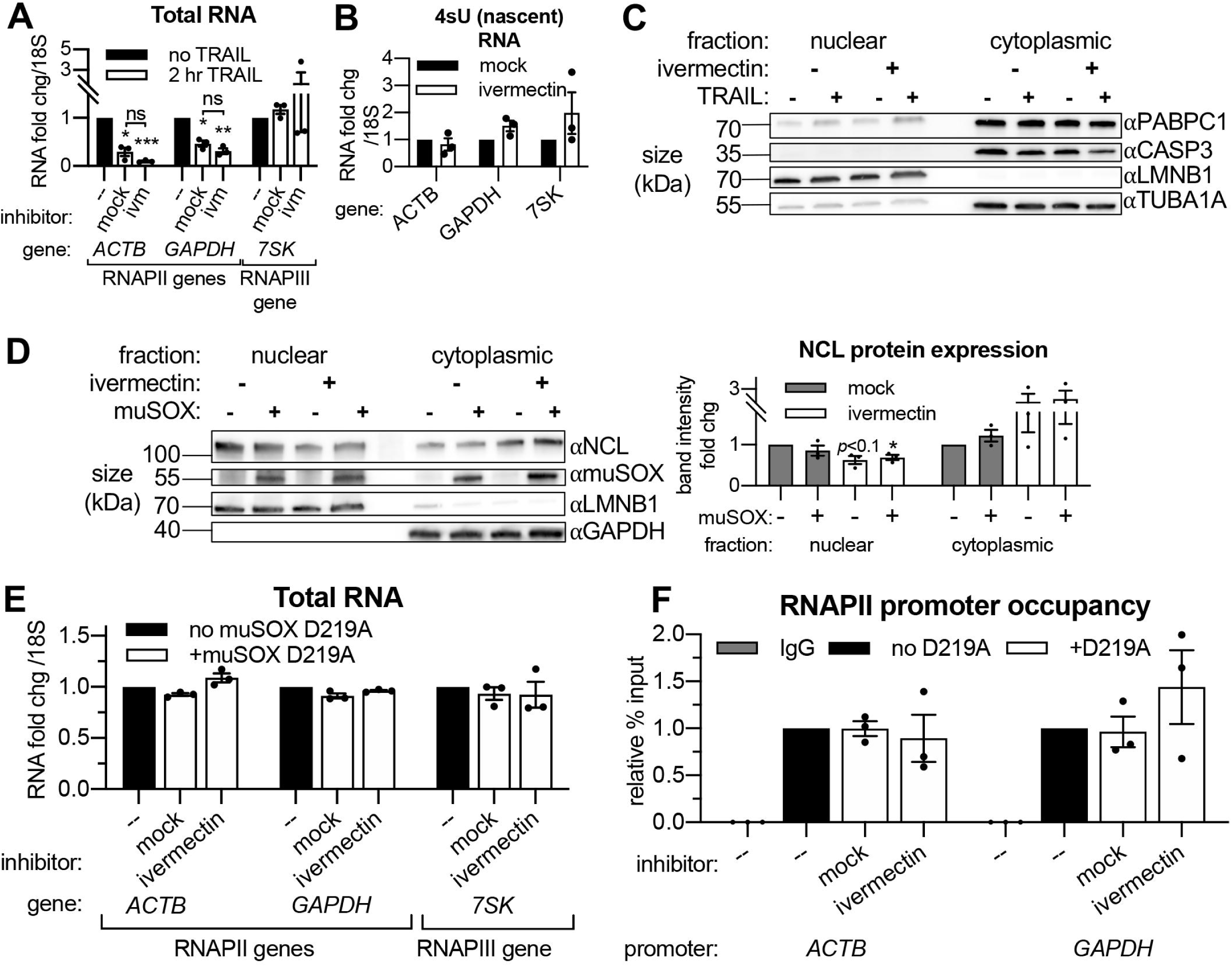
Importin α/β transport links mRNA decay and transcription. (A) Western blot showing the indicated proteins in the nuclear and cytoplasmic fractions of HCT116 cells with or without 2 hr TRAIL treatment and/or 3 hr ivermectin. Lamin B1 (LMNB1) and TUBA1A served as nuclear and cytoplasmic loading controls, repectively. Blot representative of that from 3 biological replicates. (B) Changes in total RNA levels after 2 hr TRAIL treatment in HCT116 cells pretreated with either 25 μM ivermectin or an equal volume of ethanol (“mock”) for 1 hr (*n* = 3). (C) Alternative analysis of data presented in Figure 5C, instead quantifying the fold change in nascent transcription in non-apoptotic cells in the presence or absence of 25 μM ivermectin (*n* = 3). (D) Western blot showing nuclear and cytoplasmic expression of the indicated proteins in HEK293T expressing the viral endonuclease muSOX with a 3 hr treatment of 25 μM ivermectin or equal volume of ethanol. Nuclear and cytoplasmic levels (right) of nucleolin (NCL) were quantified by band intensity in reference to a lamin B1 (LMNB1) or glyceraldehyde 3-phosphate dehydrogenase (GAPDH) loading control, respectively. (E) RT-qPCR quantification of the change in total RNA levels upon dox-inducible expression of the catalytically dead D219A mutant of muSOX with a 3 hr treatment of 25 μM ivermectin or equal volume of ethanol. (F) RNAPII occupancy at the *ACTB* and *GAPDH* promoters under cellular conditions described in (E). RNA fold change values were calculated in reference to *18S* rRNA. All bar graphs display mean +/- SEM with individual biological replicates represented as dots. Statistically significant deviation from a null hypothesis of 1 was determined using one sample t test and indicated with asterisks directly above bars, while student’s t tests were performed to compare mean fold change values for mock inhibitor or scramble treated cells to those treated with inhibitor or a targeting siRNA and indicated with brackets. \**p*<0.05, \*\**p*<0.0.1, \*\*\**p*<0.001.

## MATERIALS AND METHODS

### Lead Contact

Further information and requests for resources and reagents should be directed to and will be fulfilled by the Lead Contact, Britt Glaunsinger.

### Materials availability

The dox-inducible muSOX-expressing cell line and rabbit polyclonal anti-muSOX antibody generated in this study are both available upon request.

### Data and Code Availability

Sequencing data generated in this study are publicly available on GEO repository (accession number GSE163923).

### Cells and culture conditions

Wild-type HCT116, HEK293T, and HeLa cells (all from ATCC) were obtained from the UC Berkeley Tissue Culture Facility. Cell lines were authenticated by STR analysis and determined to be free of mycoplasma by PCR screening. HEK293Ts were made to stably express doxycycline(dox)-inducible wild-type muSOX and its catalytically-dead D219A mutant by PCR amplifying the aforementioned coding sequences from Addgene plasmids 131702 and 131704 using the muSOX F/R primers (see Key Resources Table) and InFusion cloning these fragments into the Lenti-X^TM^ Tet-One^TM^ Inducible Expression System digested with AgeI. Lentivirus was made for both constructs by transfecting 293T cells with second generation packaging plasmids and spinfected onto 293T cells at a low multiplicity of infection (MOI) as previously described (Hartenian et al., 2020). 24 hr later, 350 μg/ml zeocin was added to select for transduced cells. HEK293T and HeLa cells were grown in Dulbecco’s Modified Eagle Medium (ThermoFisher Scientific) supplemented with 10% fetal bovine serum (FBS) and 1 U/mL penicillin-streptomycin (pen-strep). HCT116 cells were maintained in McCoy’s (modified) 5A medium (ThermoFisher Scientific) with 10% FBS and 1 U/mL pen-strep. All cells were incubated at 37 °C with 5% CO_2_. Cells were maintained in culture in 10 cm^2^ plates and 1×10^6^ cells were seeded into 6 well plates for all experiments except for chromatin immunoprecipitations, in which 5×10^6^ cells were seeded into 10 cm^2^ plates, and 4sU-sequencing, in which 5×10^6^ cells were seeded into 15 cm^2^ plates.

### siRNA nucleofections

Protein knockdowns were performed using the Neon Transfection System with siRNA pools for the following targets: non-targeting control pool (scramble or scr siRNA), CASP3, CASP8, DFFB, DIS3L2, EXOSC4, PNPT1, and XRN1. For all siRNA pools, cells were nucleofected according to manufacturer protocols for HCT116 cells and immediately seeded into plates containing media supplemented with 10% FBS but lacking pen-strep to improve cell viability post-nucleofection. A 50 nM final siRNA concentration was used for pools targeting CASP3, CASP8, and DFFB (CAD), while 200 nM was used for individual knockdown of XRN1, DIS3L2, EXOSC4, and PNPT1, as well as the concurrent knockdowns of DIS3L2/EXOSC4/PNPT1 (66.7 nM each) and BAX/BAK (100 nM each). For all experiments involving siRNA knockdowns, cells were treated and/or harvested once 80-90% confluent in each plate or well, approximately 72 hr post-nucleofection, and protein knockdown was confirmed by western blot. Cell populations of siRNA-transfected cells were split into two wells or plates 24 hr pre-harvesting to allow for the direct comparison between apoptotic and non-apoptotic cells in the same genetic background.

### Apoptosis induction

100 ng/mL TNF-related apoptosis-inducing ligand (TRAIL) was used to induce rapid extrinsic apoptosis in HCT116 cells. Where indicated, HCT116 cells were pre-treated for 1 hr with 40 μM caspase Inhibitor Z-VAD-FMK (zVAD) or 25 μM ivermectin before TRAIL treatment. Extrinsic apoptosis induction was confirmed by observing CASP8 and CASP3 cleavage on western blots. CASP3/8 cleavage was quantified by disappearance in intensity of the full-sized band normalized to TUBA1A or VCL using Bio-Rad ImageLab software. Intrinsic apoptosis was induced in HeLa cells with a 4 hr treatment of 10 μM raptinal, with or without 1 hr pre-treatment with 20 μM zVAD. Induction was confirmed by evidence of cytochrome c release into the cytoplasm with cell fractionation and western blot. “Mock” treatments consisted of an equal volume of vehicle used to dissolve each reagent: TRAIL storage and dilution buffer, DMSO, and ethanol for TRAIL, zVAD, and ivermectin, respectively.

### RNA, DNA, and protein extractions

Total RNA, and protein extractions were performed according to manufacturer’s instructions after cells were harvested with Trizol^TM^ reagent. RNA pellets were dissolved in DEPC water and protein pellets were dissolved in 1% SDS overnight at 50 °C before spinning down insoluble material at 10,000 x g for 10 min.

### TUNEL Assay

Terminal deoxynucleotidyl transferase Br-dUTP nick end labeling was performed on TRAIL-treated cells using a TUNEL assay kit according to manufacturer protocol. Br-dUTP incorporation was quantified by flow cytometry, analyzing the elevated peak in FL2 fluorescence on a BD Accuri Flow Cytometry System using FlowJo analysis software.

### Quantitative reverse transcription PCR (RT-qPCR)

Extracted RNA was DNAse-treated treated, primed with random nonamers, and reverse-transcribed to cDNA with Avian Myeloblastosis Virus Reverse Transcriptase according to manufacturer protocols. Genes were quantified by RT-qPCR using iTaq Universal SYBR Master Mix and primers specific to each gene of interest. RNA fold change values were calculated in reference to *18S* or *U6* ncRNAs, as indicated on figure axes.

### Western blotting

Protein samples were quantified by Bradford assay according to manufacturer instructions. 12.5-50 μg of protein was mixed with one third volume of 4X Laemelli Sample Buffer (Bio-Rad Laboratories 1610747) and boiled at 100 °C before loading into a polyacrylamide gel alongside either the PageRuler™ or PageRuler™ Plus Prestained Protein Ladder. Proteins were separated by electrophoresis and transferred to a nitrocellulose membrane, which was then cut into sections surrounding the size of the protein of interest, allowing for multiple proteins to be quantified from one gel. Membranes were then blocked with 5% non-fat dry milk in TBST (1X Tris-buffered saline with 0.2% [v/v] Tween 20) at RT for 1 hour then incubated with relevant primary antibodies diluted with 1% milk in TBST overnight at 4 °C. All primary antibodies were applied at a 1:1000 dilution with the exception of the following (target, dilution): RPB1, 1:500; TUBA1A, 1:500; RPB2, 1:500; RPB3, 1:10000; LMNB1, 1:1000; GAPDH, 1:5000; and CYTC, 1:500. After three five min TBST washes, species-specific secondary antibodies were diluted 1:5000 with 1% milk in TBST and incubated for one hour at RT. Blots were then developed, after three additional five min TBST washes, with Clarity Western ECL Substrate for five min and imaged using a ChemiDoc^TM^ MP Imaging System (Bio-Rad Laboratories). Each membrane section was imaged and processed separately. Band intensity was quantified using Bio-Rad Image Lab software, and relative expression changes were calculated after normalizing to an α-tubulin (TUBA1A), vinculin (VCL), or lamin-B1 (LMNB1) loading control. For all blots that appear in figures, auto-contrast was applied in Image Lab for each membrane section before the resultant image was exported for publication. When appropriate, membrane sections were stripped with 25 mM glycine in 1% SDS, pH 2 and washed 2 times for 10 min with TBST before being blocked and re-probed as previously described with a primary antibody targeting a protein of similar size.

### Antibody generation

Polyclonal rabbit anti-muSOX antibody was made and purified by YenZym antibodies, LLC from recombinant muSOX protein.

### Cell Fractionations

For experiments performed in apoptotic HCT116 and HeLa cells, nuclear, cytoplasmic, and mitochondrial fractions were isolated using the Abcam Cell Fractionation Kit according to manufacturer’s instructions. Cytoplasmic and nuclear fractions of samples in experiments that did not involve apoptosis were separated using the REAP method (Suzuki, Bose, Leong-Quong, Fujita, & Riabowol, 2010). 1/5^th^ of the total cell lysate was reserved and diluted to the same volume of the cell and nuclear fractions for whole cell lysate samples. Protein was extracted from 200 μL of each fraction from both methods using Trizol^TM^ LS reagent and analyzed by western blot as described above.

### 4-thiouridine- (4sU)- pulse labeling

4sU-pulse labeling was used to measure nascent transcription concurrently with mRNA decay. 50 μM 4sU was added to cells 20 min before harvesting lysates for RNA and/or protein extraction. Labeled transcripts contained in 25 μg of total extracted RNA were biotinylated as described by Dölken (2013), using 50 μg HPDP biotin. Biotinylated RNA was conjugated to Dynabeads™ MyOne™ Streptavidin C1 magnetic beads for 1 hour in the dark, then the beads were washed four times (twice at 65 °C and twice at RT) with wash buffer (100 mM Tris-HCl, 10 mM EDTA, 1 M sodium chloride, and 0.1% Tween 20) before eluting RNA off of the beads twice with 5% BME in DEPC water. RNA was precipitated by adding 1/10^th^ volume of 3M sodium acetate and 2.5 volumes of ethanol and spun down at full speed in a 4°C benchtop centrifuge. After a 75% ethanol wash, the 4sU-labeled RNA was resuspended in 20 μL DEPC water for use in in RT-qPCR.

### Reverse transcription PCR (RT-PCR)

4sU-labeled RNA from HCT116 cells treated with 100 ng/uL TRAIL, 10 μM raptinal, or their respective mock treatments was isolated as previously described. 2 μL 4sU RNA from each sample was reverse transcribed into cDNA and amplified with primers targeting regions of the 18S rRNA and/or U6 snRNA using the QIAGEN OneStep RT-PCR Kit according to manufacturer instructions. 25 μL of each resultant PCR product was combined with 5 μL 6X DNA loading dye and loaded onto a 1% agarose (in 1X Tris-borate EDTA) electrophoresis gel stained with SYBR^TM^ Safe DNA Gel Stain alongside a DNA ladder. Gels were imaged on ChemiDoc^TM^ MP Imaging System.

### 4sU-sequencing (4sU-seq)

5×10^6^ HCT116 cells were seeded onto 15 cm^2^ plates and 24 hr later, were pre-treated with either DMSO or 20 μM zVAD, and treated with storage and dilution buffer or 100 ng/μL TRAIL for 2 hr. This process was repeated to generate 2 biological replicates. 50 μM 4sU was added to cells 20 min before harvesting. Cells were suspended in 2 mL Trizol and RNA extracted as previously described. Biotinylation and strepdavidin selection was performed on 200 μg of total RNA, scaling up the previously detailed protocol by 8X. 125 ng of 4sU-labeled RNA was used to synthesize rRNA-depleted sequencing libraries using KAPA-stranded RNA-Seq Kit with Ribo-Erase, HMR according to manufacturer’s instructions. ERCC RNA Spike-in Mix 1 was added at a 1:100 dilution to each RNA sample immediately prior to library preparation normalize read counts to RNA input across samples. Libraries were submitted for analysis on a Bioanalyzer to ensure ∼400 bp fragment lengths, then submitted for sequencing on a Nova-Seq 6000 with 100 bp paired-end reads at the QB3-Berkeley Genomics Sequencing Core.

Bioinformatics analysis was conducted using the UC Berkeley High Performance Computing Cluster. Paired end sequence FASTQ files were downloaded and checked for quality using FastQC. Reads were then trimmed of adaptors using Sickle/1.33. Reads were mapped to human reference genome hg19 and ERCC spike-in list was obtained using STAR genome aligner (Dobin et al., 2013). Differential expression upon TRAIL treatment for each gene were calculated using Cuffdiff 2 (Trapnell et al., 2013) on samples in the DMSO condition with their replicates, and on zVAD samples and their replicates. Differential expression values for each gene were normalized to ERCC spike-in controls. Normalized fold change values were calculated and analyzed using Microsoft Excel. Statistically significant >2-fold upregulated genes upon TRAIL treatment in the DMSO condition were input into PANTHER-GO Slim gene ontology analysis (Mi, Muruganujan, Ebert, Huang, & Thomas, 2019) and ChEA3 transcription factor enrichment analysis (Keenan et al., 2019).

### Chromatin immunoprecipitation (ChIP)

HCT116 cells nucleofected with the relevant siRNAs were seeded onto 10 cm^2^ plates. 72 hr post-nucleofection, cells were washed with PBS, trypsinized, washed with PBS again, and fixed in 1% formaldehyde for 2.5 min before quenching with 125 mM glycine. After an additional PBS wash, cells were lysed for 10 min at 4°C in lysis buffer (50 mM HEPES pH 7.9, 140 mM NaCl, 1 mM EDTA, 10% [v/v] glycerol, 0.5% NP40, 0.25% Triton X-100) then washed with wash buffer (10 mM Tris Cl pH 8.1, 100 mM NaCl, 1 mM EDTA pH 8.0) at 4 °C for 10 min. Cells were then suspended in 1 mL shearing buffer (50 mM Tris Cl pH 7.5, 10 mM EDTA, 0.1% [v/v] SDS) and sonicated in a Covaris S220 sonicator (Covaris, Inc.) with the following parameters: peak power, 140.0; duty factor, 5.0; cycle/burst, 200; and duration, 300 sec. Insoluble material in the shearing buffer was then spun down at full speed in a 4 °C benchtop centrifuge to yield the chromatin supernatant. 10 μg chromatin was rotated overnight at 4 °C with 2.5 μg or 4 μg ChIP-grade primary antibodies targeting RPB1 and TBP, respectively, in 500 μL dilution buffer (1.1% [v/v] Triton-X-100, 1.2mM EDTA, 6.7mM Tris-HCl pH 8.0, 167mM NaCl). 5 μL of each IP was reserved as an input sample before antibody was added. 20 μL of either Dynabeads™ Protein G or a 1:1 mixture Protein A and Protein G beads, for anti-mouse and anti-rabbit antibodies, respectively, were added to each reaction and rotated at 4 °C for at least 2 hr. The beads were then sequentially washed with low salt immune complex wash buffer (0.1% [v/v] SDS, 1% [v/v] Triton-X-100, 2 mM EDTA, 20 mM Tris-HCl pH 8.0, 150 mM NaCl), high low salt immune complex wash buffer (0.1% [v/v] SDS, 1% [v/v] Triton-X-100, 2 mM EDTA, 20 mM Tris-HCl pH 8, 500 mM NaCl), LiCl immune complex buffer (0.25 M LiCl, 1% [v/v] NP40, 1% [v/v] deoxycholic acid, 1mM EDTA, 10mM Tris-HCl pH 8.0), and TE buffer (10 mM Tris-HCl pH 8.0, 1 mM EDTA). All washes were 5 min in duration and performed at 4 °C. Beads and input samples were then suspended in 100 μL elution buffer (150 mM NaCl, 50ug/ml proteinase K) and incubated at 55 °C for 2 hr then at 65 °C for 12 hr in a thermal cycler. DNA fragments were purified with an oligonucleotide clean and concentrator kit and % input values were quantified by RT-qPCR as previously described using primers complementary to the locus of interest.

### Data visualization

Bar graphs were created using GraphPad Prism 8 software and the graphical abstract was created using the Bio-Render online platform.

### Quantification and statistical analysis

Biological replicates were defined as experiments performed separately on biologically distinct (i.e. from cells cultured at different times in different flasks or wells) samples representing identical conditions and/or time points. See figures and figure legends for the number of biological replicates performed for each experiment. Criteria for the inclusion of data was based on the performance of positive and negative controls within each experiment. No outliers were eliminated in this study. One-sample t-tests were performed on control and experimental groups for which mean fold change values were calculated, comparing these values to the null hypothesis of 1. Student’s T tests (corrected for multiple comparisons with the Holm-Sidak method when appropriate) were also performed comparing means between control and experimental groups, signified by brackets spanning the two groups being compared. All statistical analyses were performed using GraphPad Prism 8 unless otherwise noted.

### Key Resources Table

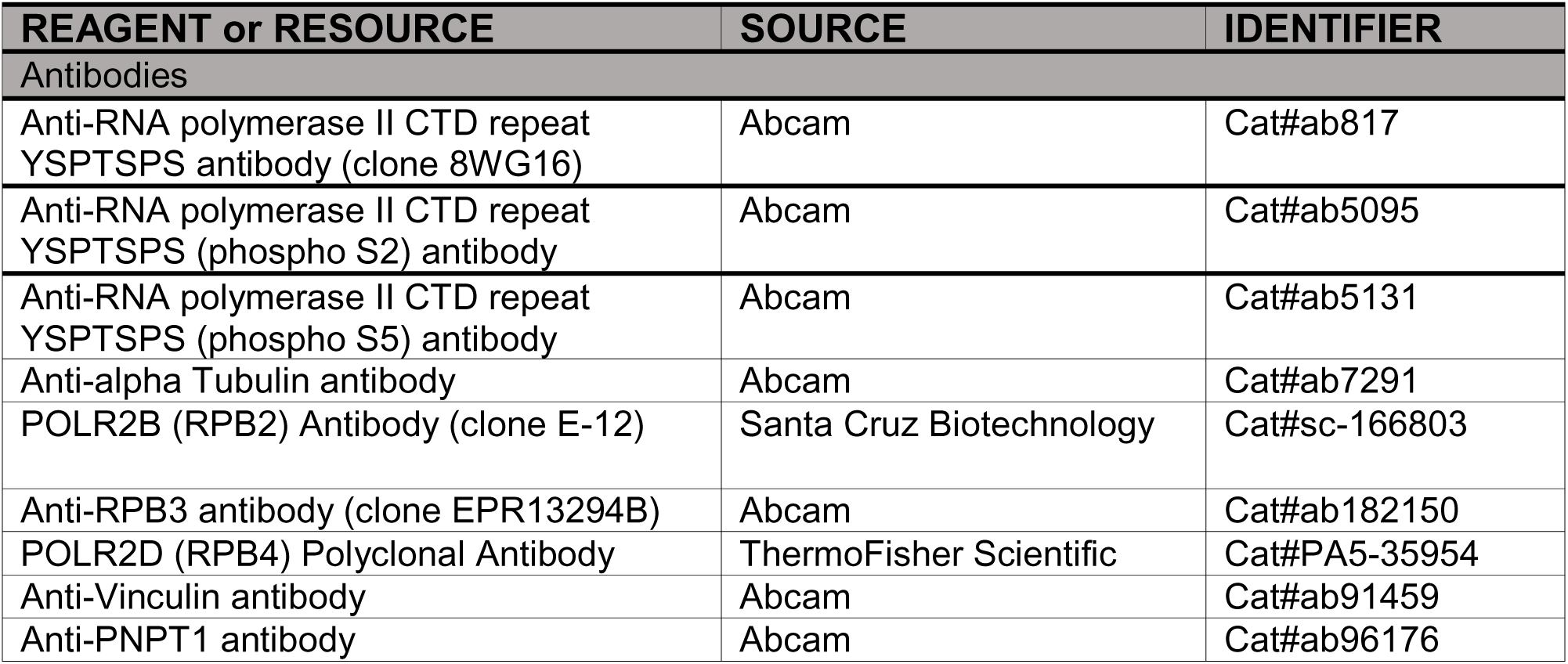
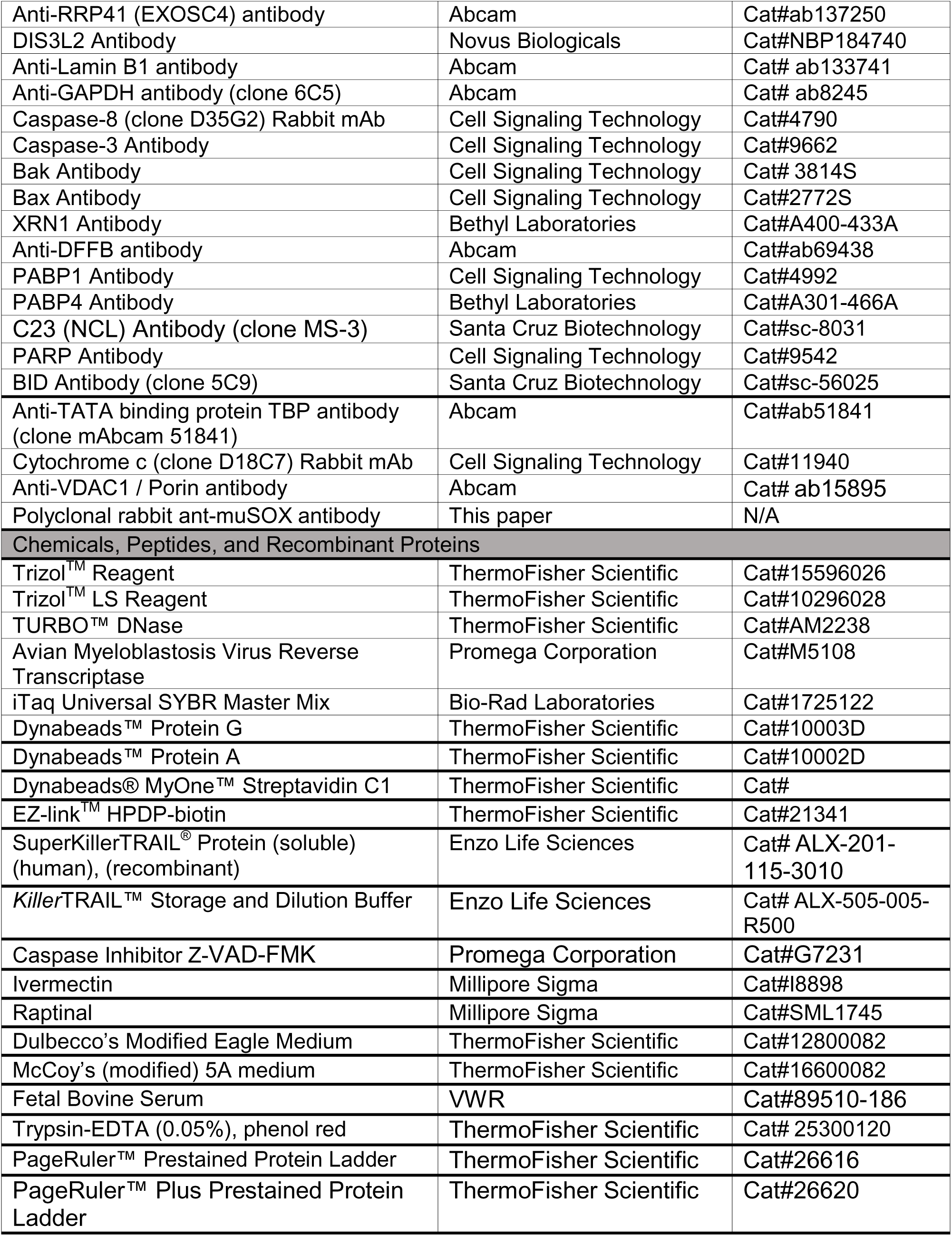
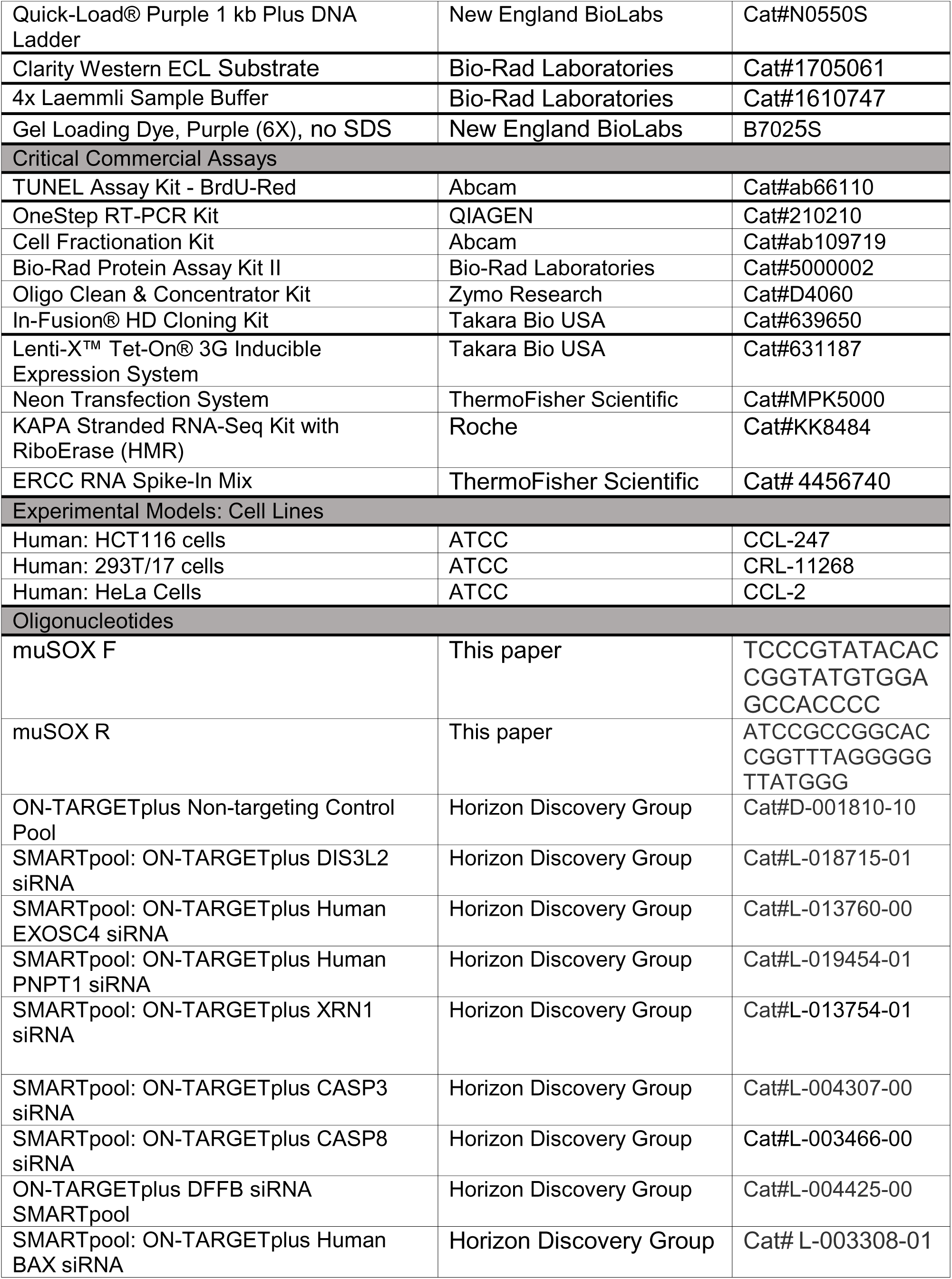
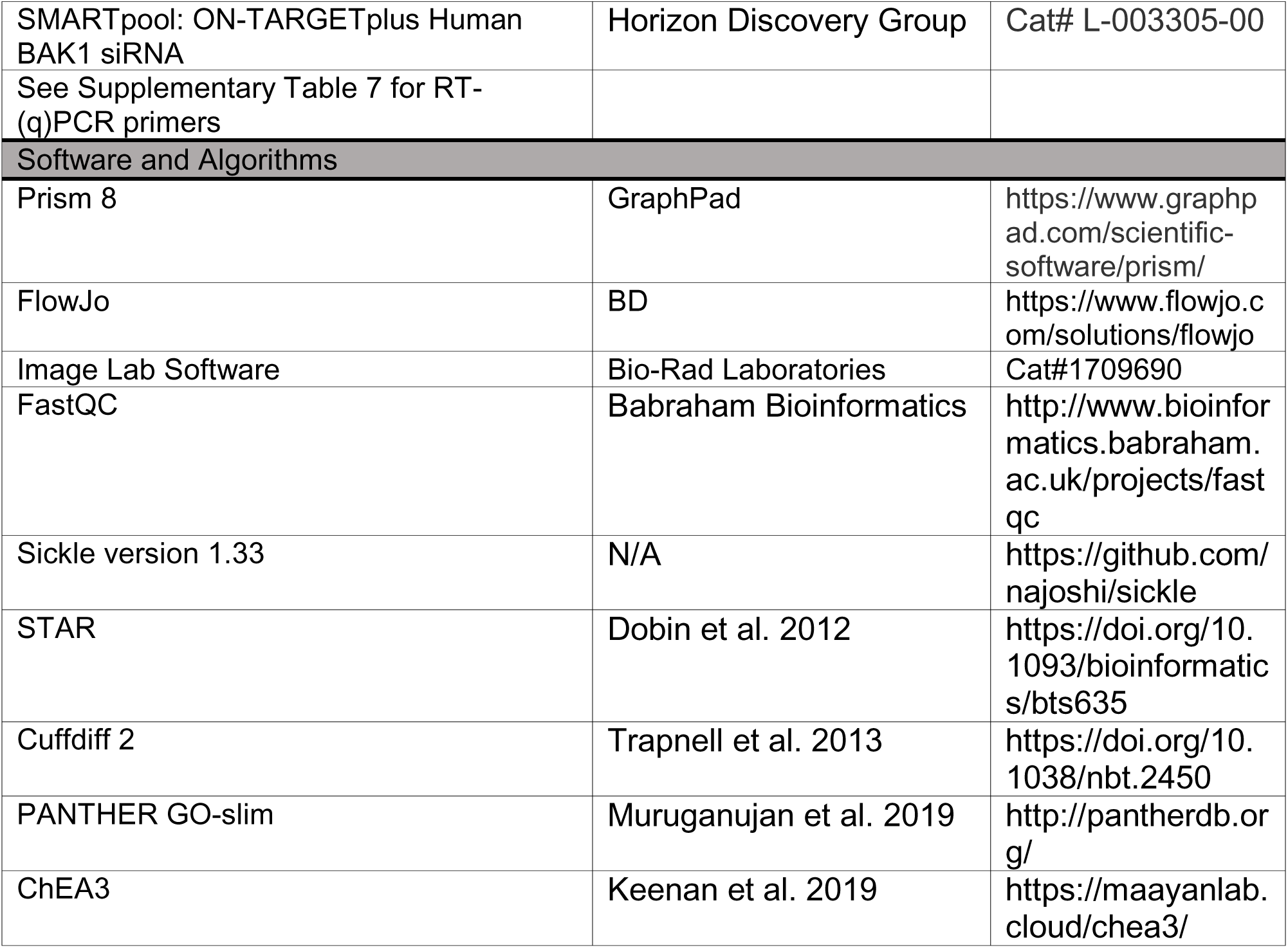

## Notes

https://www.ncbi.nlm.nih.gov/geo/query/acc.cgi?acc=GSE163923

### Competing Interest Statement

The authors have declared no competing interest.

### Summary of Updates

This version of the manuscript has been revised with additional experiments and text clarifications in response to reviewer comments.

